# ADAPT-M: A workflow for rapid, quantitative *in vitro* measurements of enriched protein libraries

**DOI:** 10.1101/2025.10.21.683815

**Authors:** Carla P. Perez, Nicole V. DelRosso, Cameron L. Noland, Udit Parekh, Christian A. Choe, Raphael R. Eguchi, Qi Wen, Polly M. Fordyce, Po-Ssu Huang

## Abstract

Protein-protein interactions underpin most cellular interactions, and engineered binders present powerful tools for probing biology and developing novel therapeutics. One bottleneck in binder generation is the scalable, quantitative characterization of these interactions. We present ADAPT-M (**A**ffinity **D**etermination by **A**daptation of **P**ro**T**ein binders for **M**icrofluidics), a streamlined workflow that connects yeast surface display (YSD) with *in vitro* affinity and kinetic measurements using the high-throughput STAMMPPING microfluidic platform. ADAPT-M quantifies *K*_d_s and dissociation kinetic parameters for hundreds of enriched protein variants in under one week without requiring hands-on protein purification. We applied ADAPT-M to a computationally designed library targeting the SARS-CoV-2 Omicron BA.1 receptor binding domain, successfully recovering and measuring *K*_d_s for most highly enriched YSD variants. Measurements correlate strongly with biolayer interferometry and yeast titration assays. ADAPT-M further enabled selection of lead candidates for structural and mutational analysis, which revealed designed paratopes were preserved despite binding to off-target epitopes. By bridging YSD screening and *in vitro* validation, ADAPT-M accelerates protein binder discovery and supports data-driven protein engineering.

## Introduction

Protein-protein interactions underlie nearly all cellular processes, from cell signaling and communication to gene expression and immune responses. Understanding and engineering these interactions is fundamental to efforts in biotechnology, synthetic biology, and therapeutic development. In particular, with the advancements seen with machine learning algorithms, which are capable of rapidly generating new designer proteins, there is an ever increasing demand for high-throughput and quantitative experimental validation methods. Display technologies such as yeast surface display^1^ (YSD) allow protein engineers to screen large libraries of designed or randomly mutated protein variants in high throughput to select for candidates binding to a target of interest. These platforms provide relative binding information across pooled variants, but not precise biophysical parameters ^1,2^. Methods such as TITE-seq^3^ and MAGMA-seq^4^ couple YSD with next generation sequencing (NGS) and fit titration curves across multiple antigen concentrations to extract quantitative affinities. While powerful, these methods can be labor- and reagent-intensive, especially for weaker or transient interactions. AlphaSeq^5^, an alternative display-based strategy, leverages synthetic yeast mating agglutination to enable library-on-library affinity profiling, using calibration to internal standards to estimate individual dissociation constants (*K*_d_s). However, this method necessitates that both protein partners be compatible with YSD, which can be limiting since large proteins, membrane-bound proteins, or proteins requiring specific glycosylation patterns may not functionally display on yeast^6^. In all cases, observed YSD enrichments convolve impacts of protein abundance (which depends on cell-specific differences in folding, degradation, and export), potential avid interactions with other surface-displayed proteins, post-translational modifications, and interaction affinities. As a result, *in vitro* characterization of hits remains essential to distinguish true binders, quantify affinities, rule out artifacts, and guide downstream development.

Traditional *in vitro* approaches for binding quantification such as surface plasmon resonance^7^ (SPR), biolayer interferometry^8,9^ (BLI), or isothermal titration calorimetry^10^ (ITC) remain low-throughput, requiring purified recombinant proteins which are often labor-intensive to produce. As a result, only a small subset of candidate sequences can be finely characterized, limiting insight into sequence-function relationships and the availability of quantitative affinity data for training predictive models. To improve the throughput, one approach has coupled cell-free expression with AlphaLISA assays^11,12^ to evaluate antigen binding and neutralization potential. This approach enables rapid, *in vitro* screening, but is unable to quantitatively measure *K*_d_s. A technology that addresses all these challenges is STAMMPPING^13^, a fluorescent-based microfluidic platform, which leverages cell free-expression and pneumatically-controlled valves to enable parallel, quantitative *K*_d_ measurements of protein-protein interactions. This method uses a C-terminal meGFP tag on immobilized binders to quantify expression levels and selectively pull down expressed proteins, thereby normalizing for sequence-dependent differences in expression and enabling high-throughput purification of protein libraries. These features make STAMMPPING a low-volume, scalable solution for directly characterizing large numbers of enriched candidates *in vitro*, making it feasible to profile the full range of diverse hits from a binder generation campaign.

Here, we introduce and demonstrate “ADAPT-M” (**<u>A</u>**ffinity **<u>D</u>**etermination by **<u>A</u>**daptation of **<u>P</u>**ro**<u>T</u>**ein binders for **<u>M</u>**icrofluidics), a workflow to adapt enriched protein binders from YSD screening for *in vitro* evaluation via STAMMPPING (**Figure 1a**). Our workflow provides a streamlined bridge between candidates enriched via *in cellulo* screening and *in vitro* measurement of affinity and kinetic constants for hundreds of putative protein binders against a shared protein target in under a week. To demonstrate the utility of the workflow, we apply ADAPT-M to a library of computationally designed binders enriched via YSD. STAMMPPING measurements of enriched binders yielded quantitative binding affinities and qualitative kinetics for 14 designs. We selected 2 candidates for detailed characterization, including paratope and epitope mapping, using STAMMPPING, YSD, and cryo-electron microscopy (cryo-EM). By enabling parallel, quantitative evaluation of a broad set of enriched binders, rather than just a select few, ADAPT-M expedites and expands the scope of protein engineering campaigns and design-build-test cycles.

**Figure 1.**
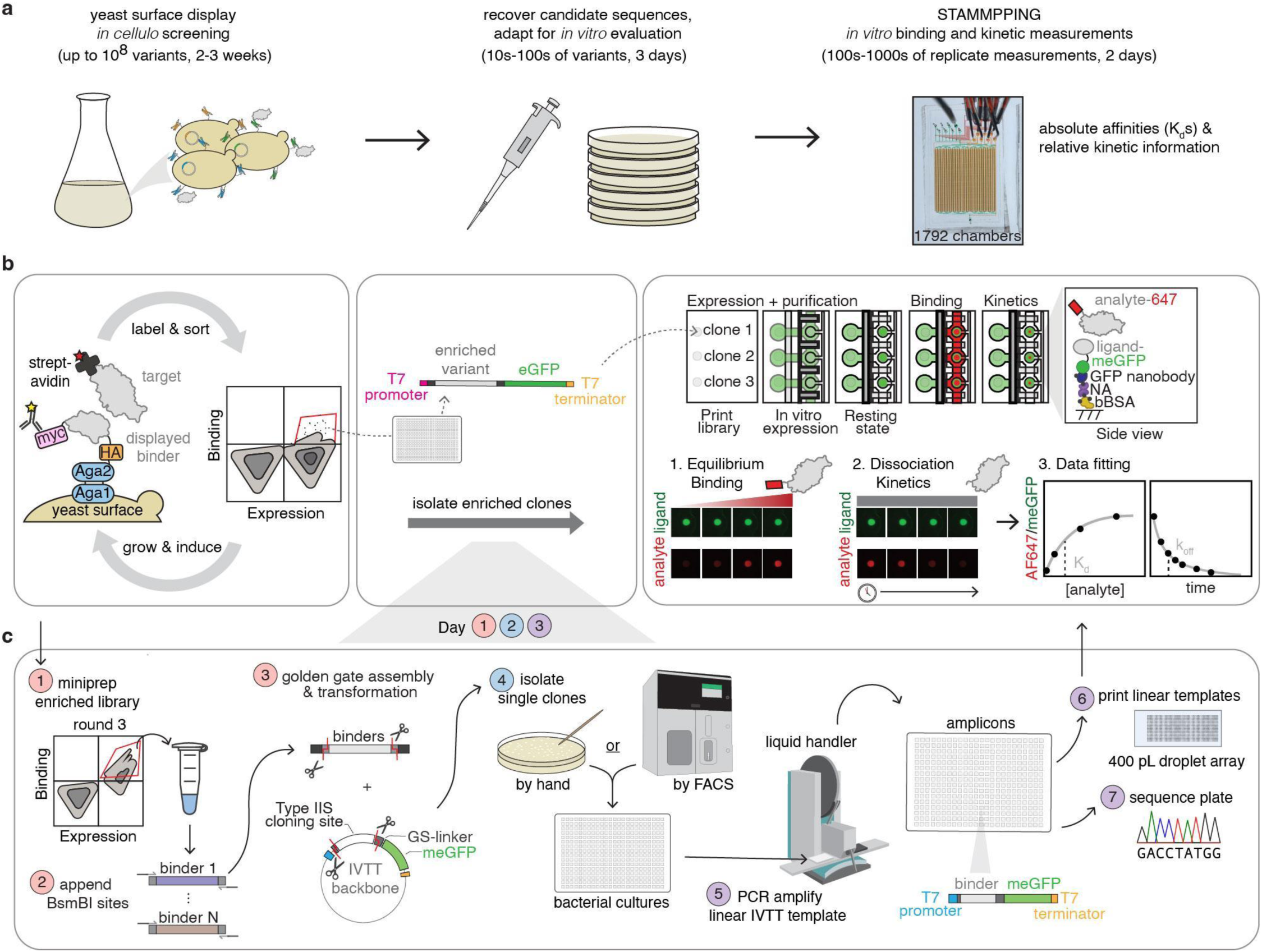
100s-1000s of affinity and kinetic measurements from enriched yeast surface display libraries in under a week. **a** Cartoon schematic illustrating an overview of the ADAPT-M workflow, starting with *in cellulo* screening in yeast, followed by recovery and adaptation of enriched DNA sequences, and concluding with *in vitro* evaluation on a microfluidic device via STAMMPPING. **b** Detailed schematic of yeast surface display (left). Designed binders are displayed on the yeast surface as Aga2 fusions with a C-terminal myc tag. Labeling with an anti-cmyc antibody permits separation of expressing and non-expressing populations during flow cytometry analysis; fluorescently-labeled target proteins discriminate between binding and non-binding populations. Binding and expressing populations are sorted, and the labeling and sorting process is repeated over multiple rounds to enrich for binders; DNA sequences from an enriched population are isolated, meGFP-tagged, adapted for cell-free expression, and directly printed for the STAMMPPING assay (right)^13^. NA=NeutrAvidin, bBSA=biotinylated bovine serum albumin. **c** Detailed schematic of ADAPT-M workflow: (1) following a few rounds of yeast surface display, the enriched population is miniprepped; (2) the binder pool is amplified and BsmBI-v2 recognition sites are appended via PCR; (3) amplicons are assembled into an cell-free expression-compatible backbone via Golden Gate Assembly and then transformed into bacteria; (4) individual clones are isolated by colony picking or by cell sorting into individual wells of a 384-well plate; (5) overnight liquid cultures are used as templates for PCR amplification of cell-free expression-compatible linear amplicons; (6) amplicons are diluted and printed in array format on a glass slide and STAMMPPING begins; and (7) the entire amplicon-containing plate is sequenced to map wells to protein variant identity.

## Results

### Overview of ADAPT-M workflow

ADAPT-M provides a pipeline to bridge pooled library screening with individual quantitative affinity measurements over 3 main stages: (1) YSD-based screening to identify candidate binders, (2) isolation and re-cloning of individual candidate binders into STAMMPPING-compatible constructs containing a C-terminal meGFP tag, and (3) STAMMPPING-based *in vitro* measurement of binding affinities (**Figure 1a**). During Stage 1, several rounds of YSD enrich a binding population with varied affinity levels (**Figure 1b**, left). Stage 2 transfers enriched variants to plasmids compatible with cell-free expression and isolates individual binder variants in unique wells of a 384-well plate (**Figure 1b**, center). Stage 3 uses STAMMPPING to express, purify, and measure up to 1,792 target interactions in parallel on a single microfluidic device (**Figure 1b**, right). This workflow makes it possible to generate hundreds to thousands of *in vitro* affinity and kinetics measurements from enriched YSD libraries in < 1 week.

#### Stage 1: Enrichment of candidate binders via YSD

Initial identification of candidate binders from libraries containing up to 1 x 10^7^ members with YSD requires 2-3 weeks (**Figure 1b**, left). During the first week, the library is transformed into yeast cells as fusions to a surface anchor protein, Aga2, permitting surface display of the library upon induction. During weeks 2 and 3, the library is induced and yeast cells are subjected to iterative rounds of fluorescence activated cell sorting (FACS) to select and enrich a target-binding population.

#### Stage 2: Preparation of linear expression templates via library parsing

Following enrichment of a binding population, the workflow shifts from pooled screening to preparation of binders for individual characterization. This involves re-cloning variants carried in a yeast surface display-optimized plasmid into a plasmid optimized for cell-free expression and fluorescent detection with STAMMPPING. These binder sequences are then physically isolated from one another over 3 days (**Figure 1c**). On Day 1, the enriched library is re-cloned into cell-free expression-compatible plasmids encoding expression of C-terminally meGFP-tagged variants via Golden Gate assembly. On Day 2, individual variant sequences are isolated by either manually picking single bacterial colonies or by FACS sorting single cells into unique wells of a 384 -well plate and incubating overnight (**Supplemental Figures 1, 2**). On Day 3, linear templates encoding cell-free expression of individual variants are amplified via bacterial colony PCR and then arrayed on glass slides to begin the STAMMPPING pipeline. Isolated clones are used directly for STAMMPPING without identifying the sequence of each clone; downstream sequencing of each template by either Sanger sequencing, short-read NGS^14,15^, or long-read sequencing^16^ allows subsequent identification of individual variants in parallel to STAMMPPING measurements.

#### Stage 3: Affinity measurements via STAMMPPING

Finally, we quantify affinities for all isolated binders via STAMMPPING, which requires 2-3 days. STAMMPPING takes place via 4 main steps: (1) polydimethylsiloxane (PDMS) devices containing pneumatic sandwich, neck, and button valves (**Supplemental Figure 3**) are aligned and bonded to glass slides bearing printed arrays (**Figure 1b**, right); (2) device surfaces are patterned with anti-eGFP nanobodies in the center of each chamber and passivated with BSA elsewhere; (3) meGFP-tagged candidate binders are expressed within each chamber and recruited to anti-eGFP nanobody-coated surfaces for on-chip purification; and (4) increasing concentrations of fluorescently-labeled target protein are introduced and mechanically ‘trapped’ at equilibrium via button valves prior to washing and imaging. Measured intensity ratios (target/binder) from images are then plotted as a function of target protein concentration and fit to quantify interaction affinities. To measure dissociation kinetics after the binding series, valves are iteratively opened to permit dissociation, closed to trap remaining bound labeled target protein, and chambers across the device are washed and imaged.

### ADAPT-M recovers enriched variants

Monobodies are small synthetic binding proteins derived from the immunoglobulin-like fibronectin domain^17^, and their compact size and lack of disulfides make them especially suitable for intracellular and therapeutic applications. Each monobody protein is composed of 7 beta strands (A-G) and 6 loops, with loops BC, DE, and FG structurally analogous to the CDR1, CDR2, and CDR3 loops of immunoglobulin heavy chain variable domains^17^. Rationally-designed monobodies typically vary these loop sequences to engage convex target surfaces, while variation of surface residues on beta strands C and D in combination with loop sequences are modified to complement more concave surfaces. Here, we applied ADAPT-M to a library of >12,000 designed monobodies targeting the severe acute respiratory syndrome coronavirus 2 (SARS-CoV-2) Omicron/BA.1 variant Receptor Binding Domain (RBD), a highly divergent variant of concern that emerged during the COVID-19 pandemic^18^. Each designed protein was 91 amino acids long (enabling library synthesis as a 300 bp nucleotide oligo pool), and designs spanned a wide range of binding orientations relative to the Omicron RBD including “loop-only” and “loop-side” recognition modes^19^ (**Figure 2a**).

**Figure 2.**
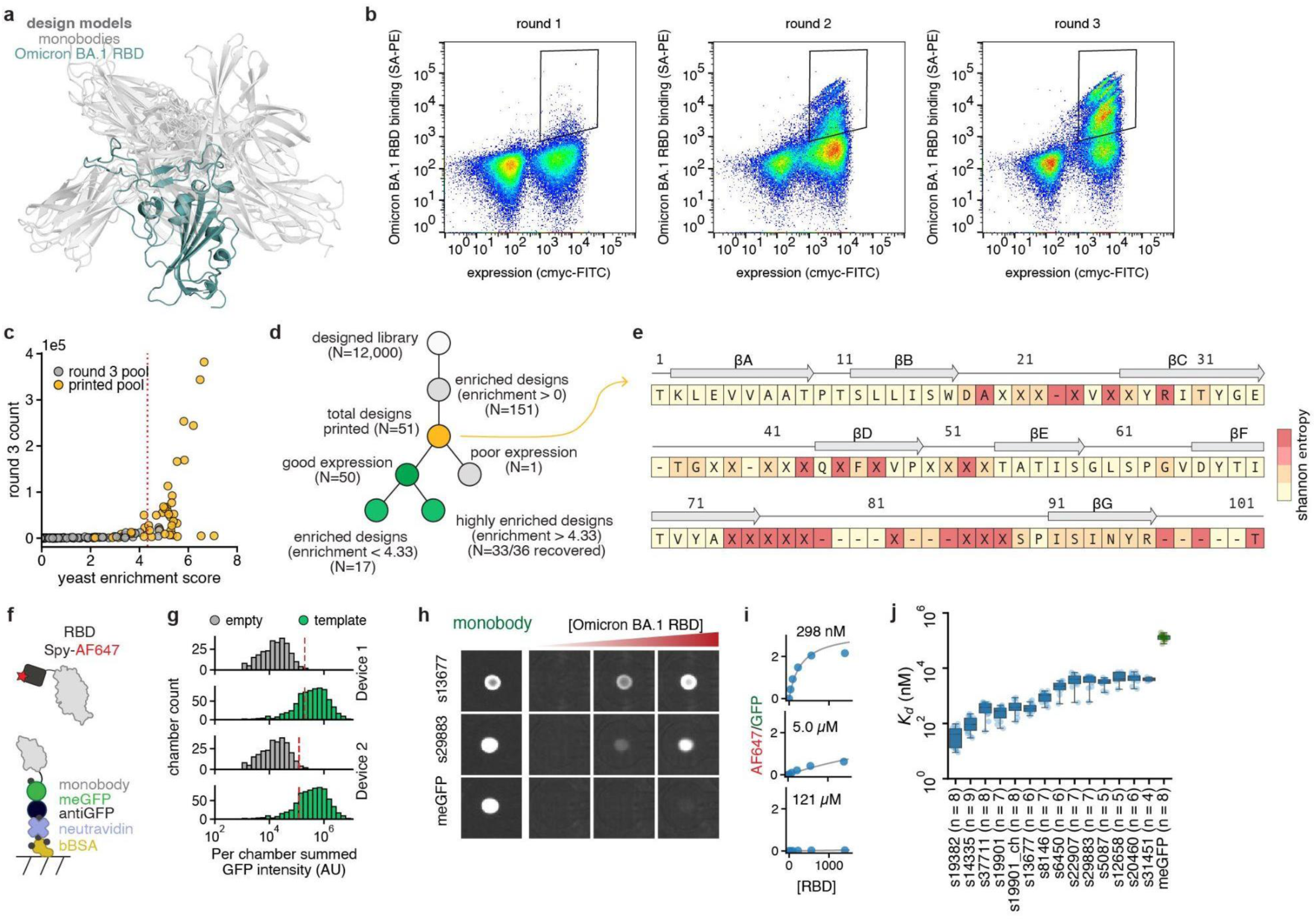
100s of affinity measurements from an enriched yeast surface display library of designed monobodies against SARS-CoV-2 Omicron/BA.1 RBD. **a** Structural models depicting 16 representative monobody designs (light grey) bound to Omicron/BA.1 RBD (teal). **b** Flow cytometry dot plots show FACS enrichment across rounds of yeast surface display selection. Approximate sort gates are outlined in black. **c** Scatter plot of round 3 counts versus enrichment scores for 151 binders (enrichment score > 0). Binders recovered for STAMMPPING measurements are shown in yellow. Red dashed line depicts the threshold for highly enriched designs. **d** Binary tree diagram summarizing outcomes for all 12,000 designed library members. **e** Schematic of sequence alignment to the monobody framework sequence used for the designed library. A secondary structure cartoon is depicted above the framework amino acid sequence. Beta strands are drawn as arrows and labeled sequentially from A to G, following their order in the amino acid sequence. Loops are represented as lines and referenced according to the strands they connect. ‘X’ denotes residue positions that were not seeded with an initial sequence and ‘–’ denotes gaps in the alignment. Amino acids are color-coded by the Shannon entropy at each amino acid position calculated across sequences printed for *in vitro* characterization. **f** Components of the functionalized device surface (which is patterned with bBSA, NeutrAvidin (NA), and a biotinylated anti-GFP nanobody that binds the meGFP-tagged monobody), and the SpyTag/Catcher-647 labeled Omicron/BA.1 RBD antigen used for measurements described here. **g** Distribution of measured intensities for empty and template-containing chambers denoting per-chamber expression for all chambers across the device. Red dotted lines represent cutoff thresholds (calculated as 10-16 standard deviations above the mean of empty chambers) used to differentiate between expressing and non-expressing chambers. **h** Example images of 3 individual reaction chambers, depicting immobilized designs after expression and purification (left) and following incubation with increasing concentrations of the Omicron RBD target (right). **i** Corresponding fits (right, solid line) to normalized Omicron RBD/monobody fluorescence intensity ratios (scatter points) as a function of free RBD concentration (**Methods**). **j** Measured *K*_d_s for enriched designs against SARS-CoV-2 Omicron/BA.1 RBD with binding above the limit of detection, variable number of replicates per design.

To identify designs that bound the target RBD, we cloned the library into a YSD vector and performed 3 rounds of FACS and outgrowth (**Figure 2b**). To maximize the dynamic range of affinities within the captured binder population, we modified 2 steps of typical binder enrichment workflows^2^: (1) we performed each round of FACS using the same high staining concentration of Omicron RBD, and (2) we maintained the same permissive FACS gating window throughout all 3 rounds of sorting. After FACS sorting, 152 variants were enriched (as assessed by computing log enrichment scores^20^ comparing variant read counts between the initial and enriched pools); 36 of these were ‘highly enriched’ (defined as having a log enrichment score > 1 standard deviation above the mean of all designs with log enrichment score > 0) (**Figure 2c**).

After selection, we used the ADAPT-M pipeline to re-clone and physically isolate enriched variants prior to STAMMPPING measurements. After individually inoculating 274 wells of a 384 well plate by hand, all inoculated wells showed successful outgrowth. Sequencing after STAMMPPING experiments revealed 51 unique monobody sequences, with a recovery rate of 92% (33/36) for the 36 highly enriched sequences (**Figure 2d**, **Table S1**). As expected, enriched monobody sequences varied most significantly in loop regions, with notable variation in the length of the FG loop of designed monobodies, as well as in the surface-exposed residues of βC and βD (**Figure 2e**). The diversity of loop sequences and conserved core observed in our selected designs align both with the sequence variability of existing monobodies in the Protein Data Bank (PDB) and our design strategy, which seeded monobody backbones with a constant framework sequence while redesigning residues at the binding interface.

### High-throughput affinity measurements

Next, we used STAMMPPING to quantify affinities between these designed monobodies and their Omicron RBD target. Fluorescence images taken after on-chip monobody expression revealed high-intensity spots corresponding to surface-immobilized meGFP-tagged proteins with intensity distributions substantially higher than those of empty chambers (**Figures 2f,g,h**); overall, 31/34 printed designs and an meGFP-only negative control expressed above the expression threshold (defined here as 10-16 standard deviations above the mean value for blank chambers) on at least 1 of 2 devices (**Figure 2g**, **Supplemental Figure 4**). After introducing Alexa 647-labeled, C-terminally SpyTag003-tagged^21^ Omicron RBD target (**Figure 2f**), additional imaging revealed concentration-dependent increases in Alexa 647/eGFP intensity ratios (which report on expression-normalized target binding), consistent with specific binding (**Figure 2h**). Fitting fractional occupancies as a function of introduced Omicron RBD concentration to a quadratic binding equation yielded *K*_d_ values that were highly replicable (Pearson R^2^=0.91, RMSLE= 0.64, **Figure 2i, Supplemental Figure 5**). *K*_d_ values spanned 3 orders of magnitude, with 14 designs exhibiting binding above the binding noise floor (as determined by a one-sided t-test with a Holm-Bonferroni correction comparing AF647 intensities of empty chambers to those of monobody-containing chambers on a per-design basis) (**Figures 2j** and **Supplemental Figures 6, 7**). Additional experiments using an orthogonal labeling approach (commercially available His-Avi-tagged Omicron RBD labeled with a fluorophore-conjugated avidin molecule, similar to the YSD experiments) also returned reproducible, high-quality binding curves (**Supplementary Figures 8**). However, while the relative categorization of strong and weak binders was consistent across experiments, the dynamic range of measured *K*_d_s for the His-Avi-tagged Omicron RBD was compressed (**Supplementary Figure 9**), making it difficult to achieve a precise one-to-one rank ordering between measurements. These compressed measurements likely result from a combination of avidity effects from using a tetrameric probe and sequence-specific avidity effects that amplify background binding. (**Supplementary Note 1, Supplementary Figures 10,11**).

### Correspondence between *in cellulo* and *in vitro* affinities

To establish the reliability of the measurements made with STAMMPPING, we selected 5 clones with varying affinities and also measured *K*_d_ values via biolayer interferometry (BLI) and yeast surface titrations^22^ (**Figure 3a**). STAMMPPING-derived *K*_d_ values correlated strongly with those obtained via label-free BLI (**Figure 3b**, **Supplementary Figure 12**, r^2^=0.83). Both measurements agreed with those derived from yeast-based titration assays (**Figure 3c**-**d**, **Supplementary Figure 13**, r^2^=0.72 and r^2^=0.8, respectively), consistent with prior reports^23^. The largest discrepancy between measurements was for design s6450, which bound more weakly by yeast-based titration than by BLI and STAMMPPING. One possible explanation for the variation in measurements is reduced epitope accessibility on the yeast surface that could be limiting the labeling efficiency of this particular design and thereby underestimating the *K*_d_.

**Figure 3.**
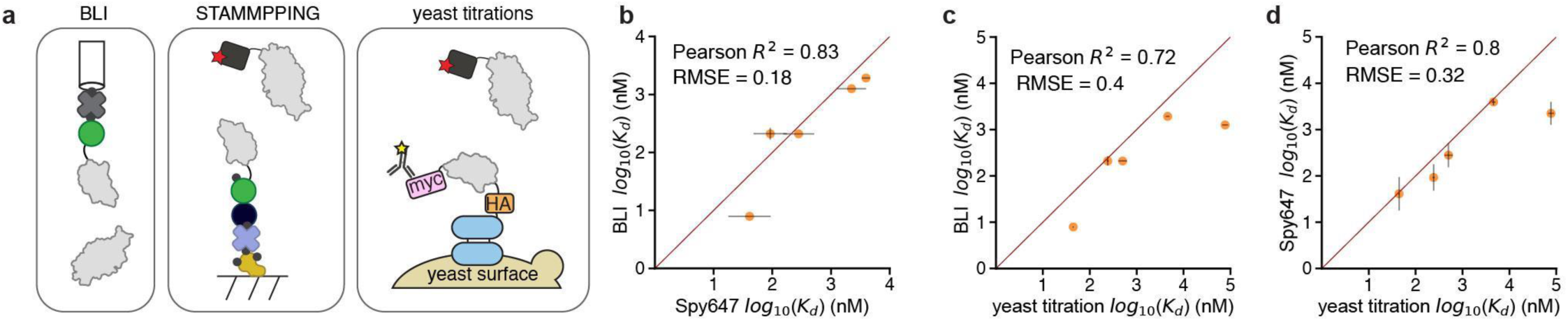
Correspondence between *in vitro* and *in cellulo* binding measurements. **a** Cartoon representation of surface chemistry for BLI measurements (left), STAMMPPING measurements (center), and yeast titration measurements (right). **b** Pairwise comparison between BLI *K*_d_ measurements (y-axis) and STAMMPPING *K*_d_ measurements (x-axis). **c** Pairwise comparison between BLI *K*_d_ measurements (y-axis) and yeast titration *K*_d_ measurements (x-axis). **d** Pairwise comparison between STAMMPPING *K*_d_ measurements (y-axis) and yeast titration *K*_d_ measurements (x-axis). Pearson R^2^ values quantify linear correlation between measurements and RMSE quantifies deviation from the 1:1 (red) line.

Overall, variants with high YSD log enrichment scores generally bound with high affinity (*i.e.* low *K*_d_) when assessed by STAMMPPING (**Supplementary Figure 14**); nevertheless, a substantial population (19/33 candidates) exhibited high enrichment via YSD and weak affinities by STAMMPPING. Observed strong binding within YSD experiments could reflect extrinsic properties inherent to the YSD environment (*e.g.* multivalent avidity effects, high local target concentrations, nonspecific binding to the yeast cell wall, and unexpected glycosylation patterns).

### High-throughput dissociation measurements

Although equilibrium binding affinities provide a useful summary of interaction strengths, they do not provide information about underlying association (*k*_on_) and dissociation (*k*_off_) kinetics that describe how rapidly complexes form and come apart. Differences in kinetic parameters can lead to distinct functional outcomes, such as enhanced efficacies from prolonged residence times^24,25^ or limited adverse effects enabled by rapid reversibility^26,27^. To begin to disentangle these processes, we monitored RBD-monobody complex dissociation over time.

Microfluidic devices similar to those used for STAMMPPING have previously been used to quantify rates of dissociation (*k*_off_s) for DNA oligonucleotides bound to surface-immobilized transcription factor proteins^28–30^. Here, we adapted this method to quantify dissociation rates for target RBD bound by surface-immobilized designed monobody binders by equilibrating binding at high target concentration, opening button valves to allow dissociation, closing them to “trap” remaining bound material, and imaging (**Figure 4a**). These dissociation measurements typically take place in the presence of a >10-fold molar excess of unlabeled target protein to prevent rebinding^31^.

**Figure 4.**
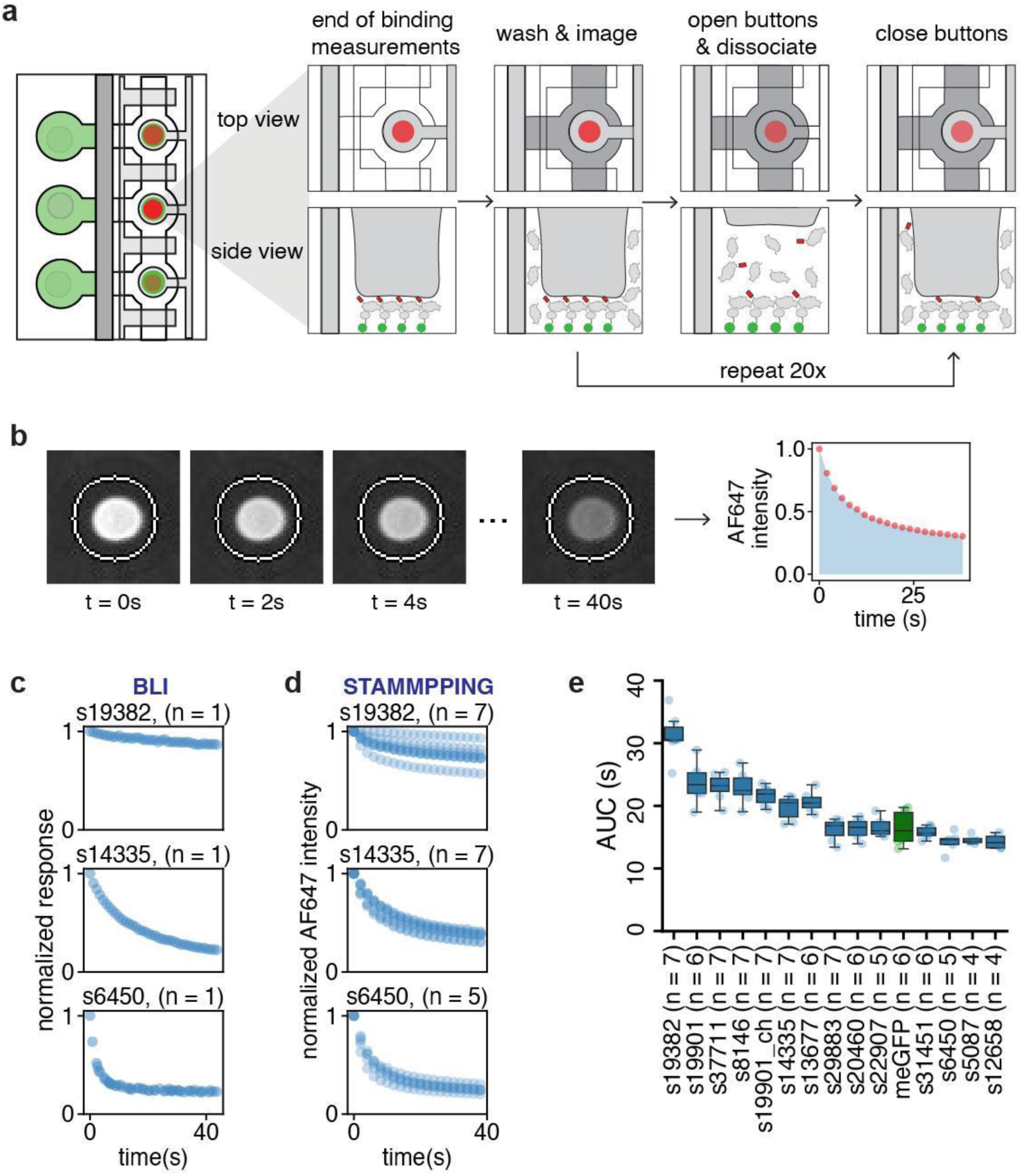
Dissociation measurements of designed monobodies via STAMMPPING. **a** Schematic of dissociation measurement protocol^28–30^ on STAMMPPING. At the end of the binding titration series, button valves are closed, and chambers are washed with an unlabeled competitor before imaging. Button valves are then opened for 2s to allow dissociation of labeled target protein, after which they are closed again to trap residual binding interactions. This cycle is repeated to monitor dissociation over time. **b** Example time course images of one reaction chamber, demonstrating dissociation of RBD over time (left). AF647 intensities are plotted against time to extract qualitative kinetic dissociation measurements (right). **c** Dissociation traces measured with BLI. **d** Dissociation traces measured with STAMMPPING. **e** Swarm plot summarizing measured AUCs for designs with binding to RBD above the limit of detection, variable number of replicates per design.

Fluorescence images of chambers revealed loss of Alexa647 signals over time, consistent with expected RBD dissociation (**Figure 4b**). However, measured STAMMPPING intensities decreased in 2 phases: a fast initial loss of RBD followed by a slower phase that did not decrease back to background levels; measurements of dissociation via BLI revealed a similar plateau (**Figure 4 c,d**). While the presence of additional molecular states complicates attempts to fit observed intensity traces and extract dissociation rates via either STAMMPPING or BLI, the traces clearly establish that RBD dissociates substantially faster from some complexes than others. To quantify these differences, we calculated the area under the curve (AUC) of fraction bound over time for each measured dissociation event. Median AUCs per design ranged from 14 - 30s (**Figure 4e**, **Supplemental Figure 15**), with binders falling into approximately 3 regimes – slow, moderate, and fast dissociators.

### Characterization of Lead Designs via Mutational and Structural Mapping

Results thus far establish that many designed monobodies sequence-specifically bind the RBD with high affinities, but do not reveal whether interactions take place at the designed interface. Here, we selected 2 designs for further characterization: s19382, the highest affinity binder with the slowest dissociation rate, and s14335, the second strongest binder, with dissociation kinetics similar to those of the other designs in this study. Notably, while s19382 ranked among the top designs (4th) by yeast log enrichment score, s14335 was only moderately enriched, ranking 23rd overall (**Supplemental Figure 14**).

#### Mutational and structural mapping of RBD epitopes

To identify the Omicron BA.1 RBD epitope bound by each design, we used YSD to express a previously-published site-saturation mutagenesis (SSM) library of Omicron BA.1 RBD^32,33^ and sorted the library separately against each binder (**Figure 5a**, **Supplemental Figures 16, 17**), reasoning that RBD mutations at the binding interface should drive loss of binding. Following sorting, we calculated positional Shannon entropies from NGS, thresholded on Shannon entropy to identify regions with mutational tolerance and on solvent accessible surface area to select surface-exposed residues, and then mapped the resulting positions onto the structure of Omicron RBD (**Figure 5b,c**)^34–36^. Both s19382 and s14335 appeared to bind a similar region on RBD, with the s19382 binding interface appearing less mutationally sensitive compared to s14335. Further, mutational scanning results suggest that both monobodies engage RBD in modes distinct from the design models (**Figure 5b**, left; **Supplemental Figure 18**).

**Figure 5.**
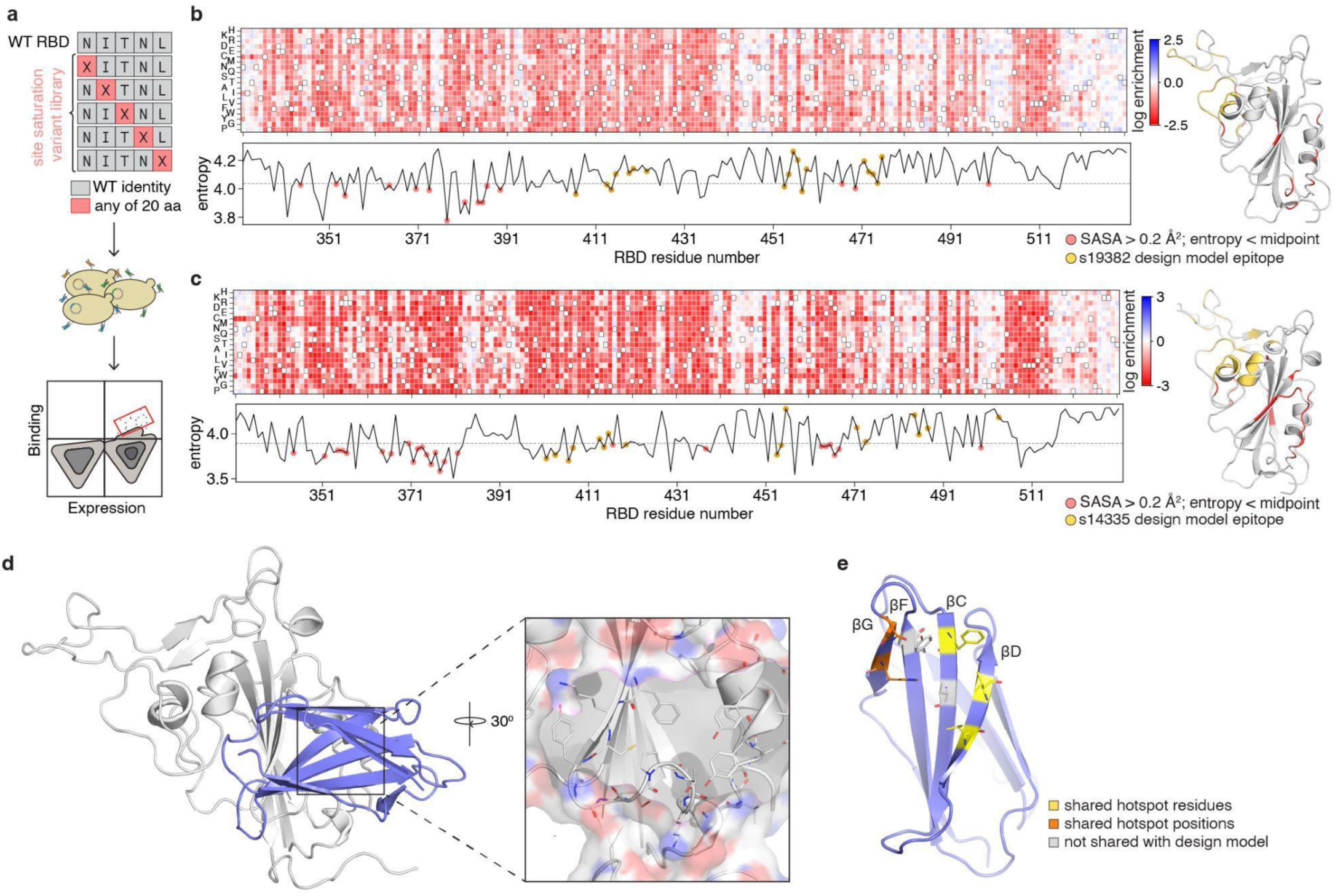
Epitope mapping of s19382 and s14335 binding via deep mutational scanning and cryo-EM. **a** Cartoon schematic of the epitope mapping strategy using YSD. A site-saturation mutagenesis library of Omicron BA.1 RBD, containing all possible single amino acid substitutions at each position (top), was expressed on the surface of yeast (center). Cells were stained with either monobody s19382 or s14335 and populations retaining binding were sorted to identify residues contributing to binding specificity (bottom). **b, c** Epitope identification for **b** s19382 and **c** s14335. Heat maps (top) derived from deep sequencing of sorted libraries, visualizing the impact of mutations on binding. Wildtype sequence outlined in black. Sequence entropy plots (bottom), with dashed black lines indicating cutoffs to define sensitive residues. Red scatter points mark residues with entropies below this threshold and solvent-accessible surface area > 0.2 Å^2^. Gold scatter points mark residues at the interface in the design model. These positions are mapped onto a structure model of RBD (right). **d** Cryo-EM structure (left) of Omicron RBD (silver) in complex with monobody s19382 (purple). A close-up view highlights the binding epitope (right). **e** Cryo-EM structure of s19382 with Rosetta-calculated hotspot residues shown as lines. Yellow indicates residues predicted as hotspots in both the design model and the experimental structure. Orange indicates paratope positions predicted as hotspots in both models, although specific contact residues differ due to beta-strand register shift. Grey indicates hotspot residues identified only in the experimental structure.

To gain insight into how these monobodies engage RBD and how their binding modes deviate from the original design models, we determined the cryo-EM structure of s19382 in complex with the Omicron RBD Spike trimer (**Supplemental Figure 19 a-e, Table S2**). Surprisingly, *ab initio* models showed that this monobody oligomerizes to facilitate Spike dimerization at its RBDs (**Supplemental Figure 19a**). In order to molecularly characterize the s19382-RBD interaction, we performed a local refinement at the dimer interface to obtain a 3.16 Å resolution reconstruction that contained two RBDs from each Spike trimer and a total of eight s19382 monobodies (**Figure 5d, Supplemental Figure 19**). Here, strong binding is facilitated by shape complementarity between the concave, β-strand–driven binding surface of s19382 and a convex protrusion on the RBD (**Supplemental Figure 20**). This interface shields a cluster of otherwise solvent-accessible aromatic residues on the RBD as well as a couple of solvent-exposed hydrophobic residues on s19382. Stabilization of the binding interface is achieved by strand-like pairing between βD of s19382 and the RBD at one end of the binding interface, and by hydrogen bonding between loop FG and the RBD at the opposite end, effectively latching s19382 onto the RBD.

This overall recognition face in the solved complex shares similarity to that of the design model (**Figure 5e**), in which s19382 binds to the RBD primarily through its FG loop and beta strands C and D. In the cryo-EM structure of s19382, packing between βA and βG is stabilized by a bulged conformation of βG and hydrogen bonding between the side chain of His80 and backbone atoms of Lys2 (**Supplemental Figure 21**). In contrast, both the design model and AlphaFold3^37^ predictions position His-80 as solvent exposed. We hypothesize that this may result from the large number of monobody structures in the PDB featuring a solvent facing Ser at this position, which could explain why neither our design model nor AlphaFold3, both trained on existing structural datasets, captured the observed packing.

The internal packing of His80 is critical to the binding mode of s19382. Packing of His80 towards the core results in a register shift in strand G as compared to the design model, altering the positions at the designed interface, particularly affecting residues on βG (**Supplemental Figure 21**). Despite the discrepancy in the binding epitope and structure of loop FG, 13 of 16 designed paratope residues on s19382 also interface with Omicron RBD in the experimental structure, with a 1.02 Å cα root-mean-square deviation (RMSD) between the monobody design model and the solved structure of s19382 (**Supplemental Figure 22**).

#### Mutational mapping of s14335 paratope residues

To identify the s14335 paratope interfacing with Omicron RBD, we used STAMMPPING to measure the effects of alanine substitutions at each surface-exposed position within the designed monobody and quantified the loss of affinity (ΔΔG) to RBD (**Figure 6a, b, Supplemental Figure 23**), reasoning again that mutations at the binding interface should drive loss of binding. Alanine scanning revealed 3 hot spot residues on s14335, Pro46, Trp48 and Thr49, with mutation of Trp48 to alanine completely ablating binding to below the limit of detection (**Figure 6c**). All 3 hotspot residues lie on loop DE, part of the designed paratope region of s14335 (**Figure 6d**), suggesting that s14335 retains at least elements of the designed paratope interface, despite binding to an alternative epitope on Omicron RBD.

**Figure 6.**
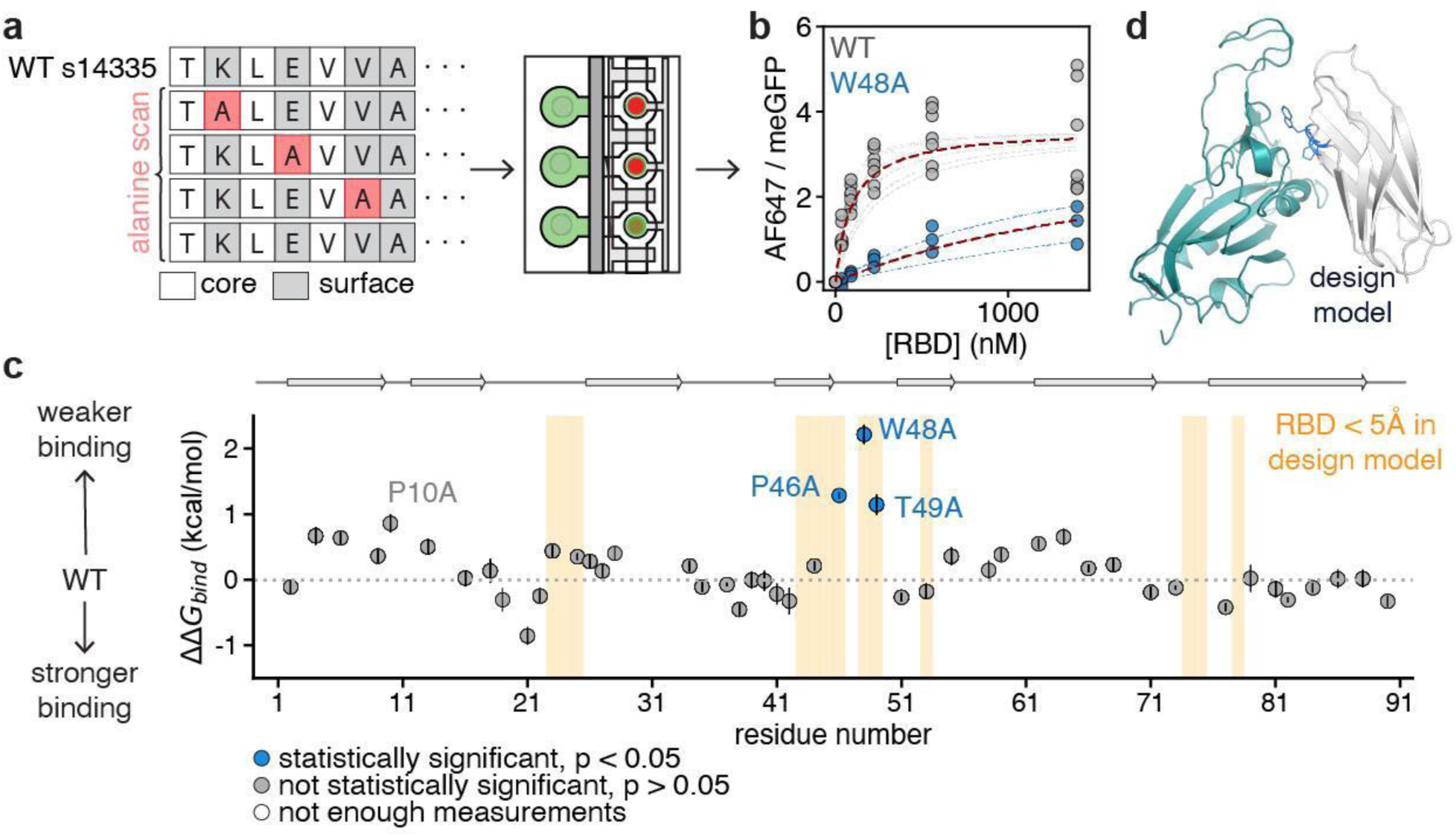
s14335 paratope mapping via STAMMPPING. **a** Cartoon schematic of the paratope mapping strategy on STAMMPPING. An alanine scan library of monobody s14335, where each residue is mutated to alanine one at a time (left), was printed for analysis via STAMMPPING (right). **b** Langmuir isotherm fits for the wildtype sequence and variant W48A. **c** Scatter plot of median per-residue contributions to the binding free energy (y-axis) plotted against residue position on s14335 (x-axis). Yellow spanning regions highlight residue positions that have an RMSD < 5Å to residues on Omicron RBD in the design model. Blue scatter points denote hot spot residues, defined as a contribution to binding of >1 kcal/mol. Grey scatter points denote residues where mutation to alanine results in a < 1 kcal/mol contribution to binding. Vertical bars on scatter points indicate standard error. **d** Design model of s14335 (grey) binding to RBD (teal). Hotspot residues identified using STAMMPPING are shown as blue lines.

## Discussion

In this study, we introduced ADAPT-M, a workflow to adapt enriched binder sequences from YSD for *in vitro* affinity determination on a microfluidic platform in under 1 week’s time. We validated the ADAPT-M workflow on a library of designed monobody binders targeting the RBD of the SARS-CoV-2 Omicron BA.1 variant. Of the 36 highly enriched candidates, 33 were successfully recovered, and binding was evaluated for a total of 50 unique design sequences with STAMMPPING. From these *in vitro* results, we selected 2 lead candidates for deeper characterization. Using YSD, cryo-EM, and STAMMPPING, we mapped the binding interfaces for both. Structural analyses confirmed that both s19382 and s14335 bind to the C4 epitope^38^ of the RBD, engaging a similar region to that recognized by the heavy chain variable domain of the Class IV binder, CR3022^39–41^. Due to this shared epitope, it is likely that these monobodies bind only in the ‘up’ RBD conformation, similar to CR3022^38^.

Through mutational and structural studies, we found both monobodies retain features consistent with the designed paratope interfaces, despite binding to an off-target epitope. This may suggest that the design metrics used here (see **Methods**), favoring large, hydrophobic interfaces, may be insufficient in evaluating alternative epitope binding modes, especially when flexible loops are involved. Moreover, the EM-resolved binding modes for these monobodies are not predictable by AlphaFold3 models, despite the high specificity and affinity observed in our experiments. The shared epitope for the 2 monobodies structurally characterized here may point to a patch on RBD that is particularly ‘sticky’ for our designed library. Indeed, library sequences that were highly enriched had a lower Rosetta ΔΔG_hydrophobic_ (*i.e.,* having a more hydrophobic interface) compared to the other sequences in the input library (**Supplemental Figure 24**). For the quantitative affinity measurements, no particular metrics, including AlphaFold3 iptm scores, appeared to have any correlation with *in vitro* binding affinity (**Supplemental Figure 25**), highlighting the limitations of current design metrics and the complexity of protein-protein recognition. These challenges also suggest why methods like ADAPT-M are critical, as they should provide rich data on many binders for further ML model improvements and can nominate particular designs for additional studies to resolve binding modes.

While we focused here on a small computationally designed library, ADAPT-M is broadly applicable to diverse protein scaffolds and library types. The workflow is not constrained by initial library size, as it leverages enrichment through YSD screening to reduce the pool to a manageable set of candidate sequences. As advances in artificial intelligence continue to transform structure-based design, ADAPT-M offers a powerful bridge to accelerate experimental validation and rapidly inform model development. The workflow is also fully compatible with mutational scanning libraries, enabling detailed studies of binding energetics and sequence - function landscapes. More broadly, we envision this workflow can be integrated into many other existing protein library characterization pipelines to extract additional biophysical insights. For example, the workflow is readily compatible with microfluidic platforms such as SPARKfold^42^, which enables high throughput measurements of protein unfolding rates via native proteolysis, providing information on protein kinetic stability.

All together, ADAPT-M provides a versatile workflow for bridging YSD screening with *in vitro* biophysical characterization, accelerating the development and functional understanding of protein binders across diverse library types.

## Acknowledgements

The authors would like to thank Zev Bryant for helpful discussions on dissociation kinetics. We would also like to thank Shohei Koide, Akiko Koide, Josh Carter, and Christopher Barnes for helpful feedback and discussion. We thank Erica Lee for help with mammalian transfections. This work was supported by NIH DP1CA290563 and NSF CAREER 2142336 awarded to P.M.F.; NIH R01GM147893 to P.-S.H. P.M.F. is a Chan Zuckerberg Biohub San Francisco Investigator. Additional support provided by Merck Research Laboratories (MRL) Scientific Engagement and Emerging Discovery Science (SEEDS) Program to P.-S.H. R.R.E was supported by the Stanford ChEM-H Chemistry/Biology Interface Predoctoral Training Program and the National Institute of General Medical Sciences of the National Institutes of Health under Award Number T32GM120007. U.P. was supported by the American Diabetes Association Postdoctoral Fellowship Program (9-22-PDFPM-05). C.A.C was supported by a Stanford Graduate Fellowship. C.P.P. was supported by a Blavatnik Family Fellowship. N.V.D. was supported by a NSF Graduate Research Fellowship DGE1656518 and an ARCS Foundation Fellowship.

## Author contributions

Project conception: CPP and P-SH

Project supervision and guidance: P-SH and PMF

ADAPT-M workflow and data collection: CPP and NVD

Monobody library design: RRE and CAC

Monobody production and testing: CPP, UP and QW

Solving CryoEM structure of s19382-RBD complex: CLN

CPP, NVD, PMF, and P-SH wrote the manuscript with contributions from all authors.

## Methods

### Computational design of SARS-CoV-2 Omicron/BA.1 RBD binders

An early version of the Sculptor^43^ algorithm was used to generate 33,095 trajectories for target binding to a defined epitope on Omicron RBD (PDB: 7tn0^44^), using a monobody generator^45^. During sequence design, a monobody framework sequence was provided, but mutations were allowed during interface design. Sequence design was completed in two stages as previously described^43^, using RosettaScripts^46^ and PyRosetta^47^ interfaces. 11,000 top ranking designs were selected by joint ranking of Rosetta metrics: ddg, interface_buried_sasa, and interface_sc. Disulfides were designed onto the 1,000 top ranking sequences and these variants were denoted by a ‘ds’ suffix.

Framework sequence: VSSVPTKLEVVAATPTSLLISWDAXXXXVXXYRITYGETGXXXXXQXFXVPXXXXTATISGLSPGVDYTITVYAX XXXXXXSPISINYRT

### Library synthesis and yeast transformation

A library of 12,000 monobody designs against the SARS-CoV-2 Omicron/BA.1 RBD was codon optimized for *Saccharomyces cerevisiae* and synthesized as a single stranded 300 nucleotide DNA oligo pool by Twist Biosciences. The oligo pool was amplified with Phusion DNA polymerase for 12-14 cycles. Long oligos were used during amplification to include upstream and downstream homology to the pCTCON2 yeast surface display vector. PCR reactions were pooled and column purified using the Qiagen PCR Purification kit and eluted into 30μL molecular grade water. 4μg of the amplified and extended library were combined with 1μg of gel purified, linearized (NheI, BamHI) pCTCON2 backbone and concentrated by a CentriVap DNA vacuum concentrator to a final volume of less than 10μL. Concentrated DNA was transformed into *Saccharomyces cerevisiae* yeast EBY100 cells following an established protocol^2^. The transformed library was grown in SDCAA media for 24 hours in a shaking incubator at 30°C. Yeast cells were then passaged into fresh SDCAA media and grown overnight. The following day, a subset of the library was stored in a 20% glycerol solution at -80°C, and the remaining subset was induced for expression.

### Yeast surface display and library screening

To induce expression, passaged yeast cells were centrifuged at 2,000g for 5 minutes and resuspended in SGCAA media at a final seeding density of 1 x 10^7^ cells/mL. Expression was induced for 18-20 hours at 30°C in a shaking incubator. To prepare yeast cells for sorting, 1-2 x 10^7^ induced yeast cells were washed three times with PBSA (PBS containing 0.5% BSA). Yeast cells were labeled with anti-c-myc fluorescein isothiocyanate (FITC, Miltenyi Biotec) (1:100) and His-Avi-tagged SARS-CoV-2 Omicron/BA.1 RBD (Acro Biosystems, SPD-C82E4) (1 μM final concentration) for 60 minutes at room temperature on a rotating platform. Yeast cells were then washed once, resuspended in PBSA, and labeled with Streptavidin, R-Phycoerythrin (SAPE, Invitrogen) (1:100) for 10-15 minutes at room temperature on a rotating platform. Following secondary labeling, yeast cells were washed once in PBSA and kept on ice as pellets. Just before sorting, yeast cells were resuspended in PBSA and passed through a 35μm mesh filter. Yeast cells were sorted using a Sony SH800S FACS instrument with 70 μm sorting chip and gated for FITC+SAPE+ signal above SAPE+ only control. Selected yeast cells were sorted into SDCAA media supplemented with 50 U/mL penicillin-streptomycin (Gibco), grown shaking overnight at 30°C, and induced for consecutive rounds of selection.

### Yeast miniprep

Yeast cells were miniprepped using a Zymoprep Yeast Plasmid Miniprep II kit (Zymo Research) with the following modifications: 1 x 10^8^ cells were used for plasmid isolation; following resuspension in 200μL Solution 1, cells underwent one to two freeze/thaw cycles prior to proceeding to step three; the amount of Zymolase used was double that recommended; and cells were incubated at 37°C for a minimum of two hours.

### NGS sample preparation and analysis

Samples were amplified using primers designed to append partial illumina adapters:

FWD:5’-ACACTCTTTCCCTACACGACGCTCTTCCGATCT

REV: 5’-GACTGGAGTTCAGACGTGTGCTCTTCCGATCT

To minimize PCR bias, a cycle number scan was set up using 8μL reaction volumes to determine the optimal number of cycles for amplification. 8μL reactions were set up as follows: 4μL 2x Q5 Mastermix (New England Biolabs), 0.4μL of forward + rev primer mix, 10μM each, 6.4ng template, water to 8μL total volume. Thermocycling conditions: 98°C, 30s → (98°C, 10s →62°C, 10s → 72°C, 15s)xN cycles → 72°C, 2 min → 4°C, Hold. The library was then amplified at this optimal cycle number in a scaled up 400μL reaction volume. The entire PCR reaction was loaded onto a 1.2% agarose gel, and gel purified using a Qiagen gel purification kit. Concentrations were determined by Qubit (Invitrogen) and submitted for paired end 250 at Azenta (South Plainfield, NJ) or paired end 300 at Neochromosome (Long Island City, NY) for NGS sequencing.

BBMerge^48^ (V39.01) was used to merge demultiplexed reads, Cutadapt^49^ (V2.6) was used to trim adapters, and Bowtie2^50^ was used to align and map trimmed reads to the library reference fasta. Custom python scripts were used to translate sequences and calculate log enrichment scores^20^ from naive and enriched library counts as well as for downstream analyses.

### PCR amplification of enriched monobody library

Primers were designed to include flanking BsmBI-v2 recognition sites. The sequence format was as follows:

forward: 5’-gatagat**CGTCTC**gctcc-[primer binding sequence]

reverse: 5’-atgctaat**CGTCTC**gatcc-[primer binding sequence]

The bolded regions represent the BsmBI-v2 recognition site. The specific full-length primers used for amplification here were:

sars_esp3i_forward: 5’-gatagatCGTCTCgctccACCAAACTAGAAGTAGTAGCC

sars_esp3i_reverse: 5’-atgctaatCGTCTCgatccCAAGTCCTCTTCAGAAATAAGC

To minimize PCR bias, a cycle number scan was set up using 8μL reaction volumes. The full-scale 200μL reaction was set up as follows: 100μL 2x Q5 Mastermix (New England Biolabs), 10μL of forward + rev primer mix (10μM each), 56ng yeast miniprep template, water to 200μL total volume. Thermocycling conditions: 98°C, 30s → (98°C, 15s → 61°C, 30s → 72°C, 20 s)x19 cycles → 72°C, 5 min → 12°C, Hold. The entire PCR reaction was loaded onto a 1.2% agarose gel, and gel purified using a Qiagen gel purification kit.

### Golden gate assembly and transformation

100ng of nvd006 (IVTT expression plasmid with C-terminal meGFP-tag, to be deposited on Addgene) was combined with 23.4ng of amplified monobody library (2x molar excess), 1μL BsmBI-v2, 1μL T4 DNA Ligase, 1μL T4 DNA Ligase buffer, and water in a final 10μL volume. A control reaction was set up as above, replacing the amplified monobody library with water. The library assembly and control reactions were thermocycled as either (42°C, 5min → (16°C, 5 min → 42°C, 5min) x 65 → 60°C, 15 min → 4°C Hold) or as (42°C, 1hr → 60°C, 5 min). Following completion of the Golden Gate reaction, tubes were kept at -20°C until transformation. 1μL of Golden Gate assembly reaction mixture was used to transform 50μL of high efficiency chemically competent *E. coli* cells (NEB, C2987H) according to the manufacturer protocol. Transformations were either plated or diluted into a 20mL volume of LB, depending on the method for single clone isolation.

### Colony picking

Following transformation, cells were plated onto 15mm LB agar plates at a density of 100-200 colonies per plate. 70μL of either LB or 2xYT medium supplemented with 50 μg/mL carbenicillin were dispensed into wells of a 384 F-bottom plate (Greiner Bio-One) using a multichannel pipette. P200 tips were used to pick single bacterial colonies and briefly introduced into each well, using a new tip in between picking and touching each tip to a single well only briefly. Plates were sealed and incubated at 37°C overnight without shaking for at least 16 hours.

### Single cell sorting of bacteria

Following transformation, cells were diluted into 20 mL LB media supplemented with kanamycin and grown overnight at 37°C in a shaking incubator to retain library uniformity^51^. In the morning, 30 μL of saturated overnight culture was used to inoculate a 3 mL culture and grown at 37°C in a shaking incubator for 2.5 -3 hours to an optical density of 0.6-1.0. About 1.5-2 hours after inoculating cultures, a Sony SH800 FACS instrument was started up, calibrated using a 70 μm chip, and droplet sorting was aligned to an F-bottom 384-well plate. 70μL of 2xYT medium supplemented with 50 μg/mL carbenicillin was dispensed into all wells of a 384 F-bottom plate (Greiner Bio-One) using a multichannel pipette. Importantly, we noticed lowered cell viability when using LB media instead of 2xYT media.

After cultures reached the target OD, 5 x 10^7^ cells (assuming 1 x 10^8^ cells/mL at OD=1) were washed once in PBSA (PBS supplemented with 0.5% BSA). Cells were resuspended in a 1mL volume and labeled with 5 μL of 1 mg/mL Propidium Iodide (PI) and incubated for 5 minutes on a tube rotator, protected from light. Following incubation, cells were washed once in PBSA and resuspended to a final cell concentration of 0.25 x 10^6^ cells/mL before being passed through a 35 μm mesh filter. Cells were loaded onto the sorter, gated by size using FSC and SSC channels, and then gated for singlets. PI-cells were sorted in single cell sorting mode at a sample pressure of 3 with an average event rate of 500-800 events per second. After sorting, plates were sealed and incubated at 37°C without shaking for at least 24 hours.

### Linear template amplification and sanger sequencing

4.224 mL of PCR mastermix was prepared in a laminar flow hood by combining reagents into a 15 mL falcon tube as follows: 844.8 μL 5x Q5 reaction buffer, 84.48 μL 10mM dNTPs, 21.2μL 100μM T7long_forward, 21.2uL 100μM T7long_reverse, 42.24μL Q4 polymerase, 3.21mL ice-cold nuclease-free water. The mastermix was kept on ice during plate preparation. The volume was calculated as 384 wells * 10uL volume/well *1.1. Primers:

T7long_forward: 5’GCGAAATTAATACGACTCACTATAGG

T7long_reverse: 5’CTTTCAGCAAAAAACCCCTCAAG

A mosquito LV liquid handler (SPT Labtech) was used to aspirate and dispense 50nL of overnight liquid cultures and dispense into individual wells of a 384-well PCR plate (BioRad). A mantis microfluidic liquid dispenser (Formulatrix) was used to dispense 10μL PCR mastermix into each well. Plates were tightly sealed, lightly vortexed, and centrifuged before being placed into a thermocycler with a 384 -well reaction module (BioRad). Thermocycling conditions: 98°C, 3 min → (98°C, 10s → 63°C, 20s → 72°C, 50 s)x30 cycles → 72°C, 5 min → 4°C, Hold.

2-4uL of 384-well plate samples were submitted for Sanger sequencing to Elim Biopharmaceuticals (Hayward, CA). Sequencing chromatograms were manually inspected and used to map wells back to their corresponding design sequence in the library. Only STAMMPPING chambers corresponding to wells containing a single design sequence were selected for downstream analysis.

### Plasmid template preparation

For validation experiments with Spy647-labelled RBD, wells containing designs to be validated were PCR amplified with primers appending BsmBI-v2 overhangs to the protein coding sequence as previously described, and reactions were PCR purified.

4μL-scale Golden Gate assembly reactions were set up in a 96-well plate (0.4μL BsmBI-v2, 0.4μL 10x T4 DNA Ligase buffer, 0.4μL T4 DNA Ligase, 0.8μL 10ng/μL insert, 34ng nvd006 template, water to 4μL). Reactions were thermocycled overnight as follows: 42°C, 5min → (16°C, 5 min → 42°C, 5min) x 65 → 60°C, 15 min → 4°C Hold. 2μL of each Golden Gate reaction were transferred into a fresh 96-well plate and a 20uL volume of DH5alpha cells (Zymo) was added to each well for transformation. Following transformation, a 245mm agar plate was divided into an appropriate number of grids, and 4uL of each transformation was plated onto individual grids. Plates were grown overnight at 30°C. A single colony was cultured and miniprepped using a Qiagen miniprep kit. Plasmids were sequence validated.

### s14335 alanine scan library template preparation

A custom python script was used to identify residues with a SASA > 0.2 Å^2^ in the design model and generate a fasta sequence file with the mutated DNA sequence. The residue list was inspected via visualization in Pymol and updated as necessary. BsmBI-v2 recognition sites were appended to both ends of the coding sequence. The GCG alanine codon was selected for all alanine mutations. Sequences were ordered from IDT as e-blocks in a 96-well plate. Plasmids were prepared as in the section above. Flanking sites:

5’ flank: gatagat**CGTCTC**gctcc

3’ flank: ggatc**GAGACG**attagcat

### Preparation of epoxysilane-coated slides

Plain 25×75mm glass slides (Corning) were functionalized with epoxysilane (based on ref^52^). First, slides were cleaned in an 80°C heated bath of hydrogen peroxide and ammonium hydroxide for 30 min, rinsed, and blow-dried with nitrogen. Next, epoxysilane was deposited by submerging slides in toluene doped with 3 -glycidyloxypropyl-trimethoxymethylsilane (3-GPS, Millipore-Sigma) for 20 min. Slides were then rinsed in toluene and dried by baking at 120°C for 30 minutes.

### Printing DNA libraries onto slides

5-10 µL of DNA templates were transferred to a 384-well plate, and resuspended in 2x “print buffer” (20 mg/mL UltraPure BSA, 24 mg/mL trehalose dihydrate, 100 mM NaCl). Templates were randomly assigned positions within a 32 x 56 array, with controls containing only print buffer distributed throughout. Two 300 -360 pL spots were arrayed on epoxysilane-coated glass slides using a SciFLEXARRAYER S3 (Scienion) outfitted with a PDC-70 Piezo Dispense Capillary nozzle with Type 1 coating. Between every sample, the nozzle was washed to ensure no cross-contamination. Fabricated devices were then aligned on top of this array by eye, placing one spot per DNA chamber using a stereoscope. Devices were baked on a hot plate (Torrey Pines) for 4 hours at 80°C.

### Mold and device fabrication

Two-layer microfluidic devices were cast from previously fabricated 15 µm tall flow and control molding masters^13^ using polydimethylsiloxane (PDMS) polymer (Momentive, RTV615). Control layers were generated by pouring a mixed 1:5 ratio of cross-linker to base (THINKY 3 minutes at 2000 rpm) onto the molds, degassing to remove all air in a vacuum chamber for 45 minutes, and baking for 60 minutes at 80°C. Flow layers were generated by spin-coating a 1:20 ratio of mixed cross-linker to base onto the molds at 500 rpm for 10 seconds followed by 1800 rpm for 75 seconds. Layers were relaxed on a flat surface for 5-20 minutes at room temperature prior to baking at 80°C for 40 minutes. Cut and punched device control layers were then aligned to flow layers using a stereoscope. Aligned devices were then baked for 50 minutes at 80°C and excised from the molds using a scalpel.

### Microscopy instrumentation

Chip imaging was performed using a Nikon Ti-2 Microscope equipped with motorized XY-stage (Applied Scientific Instruments, MS-2000 XY stage), LED light source (Lumencor SOLA SE Light Engine), 4x Objective (CFI Plan Apo 4X NA 0.20, Nikon), and CMOS camera (Andor Zyla 4.2). The Nikon Ti-2 core itself was equipped with an automated Z-axis, turret aperture, and filter turret, containing pre-arranged filter cubes for EGFP (Semrock, GFP-4050B) and Cy5 (Semrock, Cy5-5070A). All imaging was performed using 2×2 pixel binning with exposure times as follows: EGFP: 100 ms and 500 ms, Cy5: 50 ms, 100 ms, 500 ms, 1000 ms, and 3000 ms.

### STAMMPPING device operation and experimental pipeline

Microfluidic valves were operated via a custom-built open-source WAGO-controlled pneumatic manifold^53^. Automated control of pneumatic valves and microscopy was run through custom Python scripts (RunPack, AcqPack) as described previously^54^. Briefly, standard STAMMPPING experiments consist of the following steps: (1) pattern button-valve surfaces for bait immobilization, (2) express bait-meGFP constructs from DNA templates on chip, (3) iteratively introduce labeled prey protein, allow interactions to come to equilibrium, trap bound material using ‘button’ valves, wash away unbound material, and image in both the eGFP and Cy5 channels to measure binding, and (4) open the ‘button’ valves for a defined period of time to allow dissociation of prey protein before closing and imaging in both eGFP and Cy5 channels to measure dissociation kinetics. Steps 1 and 2 were carried out largely as described previously^13,54,55^, with the following modifications. For surface patterning, we used a biotinylated anti-GFP nanobody (Proteintech, gtb-250). We also optimized on-chip expression for monobody-meGFP constructs such that expression conditions varied from 25-37°C for 2-6 hrs. Following expression and binding of monobody-meGFP to anti-GFP-patterned buttons, device chambers were washed in TrypLE and regions surrounding the protected buttons were re-patterned with biotinylated BSA or Neutravidin (**Supplementary Note 1**). For step 3, dye-labeled proteins made off-chip were prepared as serial dilutions in an Assay Buffer of 20 mM KH_2_PO_4_, 200 mM KCl, 5 mg/ml UltraPure BSA (Thermo Fisher), 0.05% v/v Tween-20 (Thermo Fisher) in UltraPure distilled water. On-chip iterative measurements of binding over multiple concentrations were carried out via scripts based on those used previously^13^.

### STAMMPPING image processing

Raw 7×7 image rasters of fluorescence images covering the full chip area were flat-field corrected using manual flat-field images or an implementation of BaSiCpy^56^ and stitched into single full-chip images using ImageStitcher, an in-house publicly-available software package (https://github.com/FordyceLab/ImageStitcher). These stitched images were processed using ProcessingPack, an in-house publicly-available software package (https://github.com/FordyceLab/ProcessingPack). Briefly, circle-finding algorithms identified fluorescent spots under chamber “button” valves and per-spot metrics were mapped to Scienion print array position. Downstream analysis primarily utilized background-subtracted summed intensities of GFP and Cy5 per-spot fluorescence.

### STAMMPPING quality control

Prior to fitting curves to determine binding affinities, we performed several quality control steps. First, we filtered out any chambers with GFP expression levels above the threshold set by considering blank chambers (mean + ten or more standard deviations). Second, we manually culled chambers that had visible dust or flow issues. Third, we computed a linear regression between measured Cy5 intensities of soluble prey in each chamber and the estimated input concentration and required r^2^>0.9 (ensuring that soluble prey was available for binding as expected, **Supplemental Figure 26**).

### Determining fractional occupancies

Given the observation of decreasing meGFP intensity over the course of binding measurements and increasing meGFP intensity over the course of kinetics measurements (**Supplemental Figure 27**), we suspected Förster Resonance Energy Transfer (FRET) between meGFP and AF647 fluorophores. FRET would artificially increase cy5/meGFP ratios at increasing Omicron RBD concentrations, to differing degrees for different monobody designs, owing to differing binding modes and strengths. To circumvent this, we use meGFP intensities measured at step 0, prior to any Omicron RBD introduction, to calculate response ratios at each step as AF647/meGFP_step0_, thereby making the assumption that meGFP is not lost due to dissociation or button “shearing” over the course of the binding experiment. For kinetics measurements, we make this same assumption, and instead fit data to per-chamber summed AF647 intensities rather than AF647/meGFP ratios.

### Determination of STAMMPPING equilibrium binding constants

All *K_d_*s were estimated by global fitting of a quadratic solution to 1:1 binding equilibrium^57^ (Equn. 1) to concentration-dependent data. Here, the concentration of immobilized monobody in each chamber, was determined by dividing the summed, background subtracted eGFP intensity for each chamber by an empirically-calculated slope (here, slope = 43,000 a.u./nM^13^), relating eGFP intensity to immobilized eGFP concentration, as has been demonstrated previously^54^.

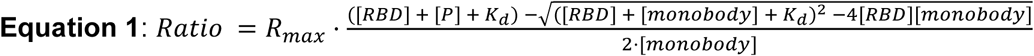

In cases where ligand (putative binder) depletion is negligible, the data can be accurately fit using the standard Langmuir isotherm (Equn. 2) instead.

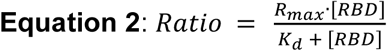

For binding measurements, we used global fits assuming that all binder/target interactions bind with the same stoichiometry to accurately determine *K*_d_s even for interactions that did not saturate at the highest concentrations tested. First, we estimated the saturation point for a given experiment by fitting the top binders (binders with a Cy5/GFP Ratio above a set threshold set to identify most-saturated binders) to Equn. 1 with *K_d_* and *R_max_* as free parameters. The mean *R_max,top_binders_* was then used as a global *R_max_* to re-fit all remaining RBD-monobody interactions with *K_d_* as a free parameter. From these fits, we calculated the median *K_d_* value across replicates for each binder/target pair. In all cases we remove outliers (> 1.5 times the interquartile region (IQR) and report median *K*_d_s. All data fitting was done in custom Python scripts using open-source lmfit and scipy packages^58,59^.

### Qualitative measurement of dissociation kinetics

Qualitative dissociation kinetics were estimated by calculating the AUC from time-course plots of summed AF647 fluorescence intensities across all regions of interest, over the first 40s of the dissociation measurement. AUC was computed by numerical integration using the composite trapezoidal rule implemented in NumPy^60^ (numpy.trapezoid).

### Determination of limit of detection

Due to the higher binding to “empty” chambers as compared to meGFP-only containing chambers, we used “empty” chambers to establish the limit of detection (**Supplemental Figure 6**). To identify designs with binding above this limit of detection, we applied a one-sided t-test with a Holm-Bonferroni correction to compare the cy5 intensities of all empty chambers to the cy5 intensities of monobody-containing chambers, on a per-monobody basis. Monobodies with p < 0.05 were denoted as binding above the limit of detection and those with p > 0.05 were categorized as having binding below this limit. Importantly, using empty chambers to establish the limit of detection provides a strict binding threshold, as meGFP-only containing chambers consistently exhibited weaker nonspecific binding than empty chambers.

### ΔΔG estimation

ΔΔG values were calculated by subtracting the wildtype sequence ΔG (Equn. 3) from each alanine variant ΔG, using a lower limit *K*_d_ when applicable. ΔΔGs were calculated per-device, and the median ΔΔG is reported.

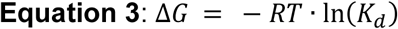

### SARS-CoV-2 Omicron/BA.1 protein expression and purification

Constructs encoding the SARS-CoV-2 Omicron BA.1 RBD (residues R328-L533 of GenBank QHD43416.1 sequence (G339D, S371L, S373P, S375F, K417N, N440K, G446S, S477N, T478K, E484A, Q493R, G496S, Q498R, N501Y, Y505H)) and either a C-terminal 3xFLAG tag or C-terminal SpyTag003 were transiently transfected into Expi293F cells (Gibco). After addition of enhancers, cells were grown at 30 °C and 8% CO_2_. Supernatants were harvested 4-5 days after transfection, and supplemented with 0.02% sodium azide (1000x) and 0.2M PMSF (100x) before being spun down at 3,500g for 15 minutes and vacuum-filtered using a 0.22um mesh filter. Filtered supernatants were diluted 2-fold in ice-cold PBS before being loaded for nickel affinity chromatography at 4°C. RBD elutions were further purified by size-exclusion chromatography (Superdex 75 or 200 10/300 GL columns) at 4°C.

### SARS-CoV-2 Omicron/BA.1 fluorescent labeling (SpyTag-SpyCatcher-647)

2mg of SpyCatcher003Cys (Bio-Rad) was resuspended in PBS in a final volume of 497.5μL. To reduce disulfides, 2.5μL of a freshly prepared 1M DTT stock was added to reach a final concentration of 5mM DTT, and the reaction was incubated at room temperature for 1 hr. Meanwhile, a Superdex 75 10/300 GL column was equilibrated with degassed PBS. After incubation, the reduced SpyCatcher003Cys was immediately injected onto the column, and monomeric fractions were collected. To prevent the reformation of disulfides, conjugation with maleimide was initiated immediately.

Monomer fractions were pooled in a 15mL falcon tube with a mini stir bar. From Nanodrop A280 measurements, typical recovery following size exclusion was 1-1.2 mg of SpyCatcher003Cys. A 10mM stock of AF647-C2-maleimide (ThermoFisher) was prepared by dissolving the full 1mg vial in 80 μL of solvent, providing a 10-20 fold molar excess relative to the protein. All 80μL of the dye stock were added dropwise to the reduced SpyCatcher003Cys, while stirring. The reaction was protected from light and allowed to proceed under vacuum at room temperature for 2 hours. Following conjugation, the protein was buffer exchanged via a Superdex 75 10/300 GL column. Labeling efficiency was calculated according to the manufacturer’s protocol, and typically exceeded 97%. The labeled protein was then concentrated using a 3 kDa MWCO Amicon filter (Millipore Sigma) to a working concentration of 80 μM.

After confirming successful labeling, SpyCatcher003-AF647 was reacted with Omicron BA.1-SpyTag003 at a 1.2:1 ratio, at a final 40μM concentration, overnight at 4°C. The next day, the conjugated protein was purified using a Superdex 200 10/300 GL Increase column equilibrated with ice-cold PBS. Fractions were analyzed by gel electrophoresis to confirm conjugation, pooled, and aliquoted. Aliquots were stored at 4°C for immediate use or -80°C for long-term storage.

### Monobody protein purification and biotinylation

meGFP-tagged monobody sequences were amplified from the print plate using primers appending 5’ and 3’ overhangs to pET24-a(+) with and without an Avi-tag at the C-terminus of the fusion protein. Expression plasmids were assembled using gibson assembly in DH5alpha cells (Zymo research), miniprepped, sequence verified, and transformed into BL21(DE3) *E.coli* cells (Zymo Research). A single colony was used to inoculate 5 mL 2xYT cultures supplemented with kanamycin and grown in a shaking incubator at 37°C for at least 5 hours. Cultures were then diluted in 500 mL 2xYT and grown until the optical density reached 0.6-1.0. Protein expression was induced by addition of isopropyl β-D-thiogalactopyranoside (IPTG) to a final 1mM concentration and cultures were grown at 16°C for 16-20 hours. Following induction, cells were pelleted and resuspended in 30 mL of phosphate buffered saline (PBS), pH 7.4 and frozen at -20°C until purification.

Pellets were thawed at room temperature and 4.2mL of 4M NaCl and 600μL of 50 mM phenylmethylsulfonyl fluoride (PMSF) were added. Thawed pellets were sonicated, clarified by centrifugation, and purified by nickel affinity chromatography followed by size-exclusion chromatography (Superdex 75 Increase 10/300 GL or Superdex 200 Increase 10/300 GL, GE Healthcare) into either 50 mM Bicine, 100 mM NaCl, pH 8.3 buffer (Avi-tagged) or PBS (non-Avi-tagged). Elutions were characterized by SDS-PAGE and concentrations were determined by absorbance at 280 nm measured with a NanoDrop8000 spectrophotometer (Thermo Fisher Scientific) using predicted extinction coefficients.

Avi-tagged proteins were biotinylated using a BirA biotin-protein ligase standard reaction kit (Avidity), and the reaction was incubated overnight at 4°C followed by one hour at room temperature. Proteins were then buffer exchanged into PBS, pH 7.4 buffer via size-exclusion chromatography to remove unreacted biotin. The extent of biotinylation was assessed by streptavidin gel-shift analysis^61^.

### Biolayer interferometry

BLI experiments were performed on an Octet RED96 system (Forté Bio). Octet streptavidin (SA) biosensors (Sartorius) were rehydrated in a 1x kinetics buffer (Sartorius) for at least 30 minutes at room temperature. Biotinylated meGFP-tagged monobodies diluted in 1x kinetics buffer to a concentration of 0.8 ug/mL. FLAG-tagged Omicron RBD stocks were diluted first in PBS to 4-5x the target concentration and then diluted 4-5-fold into 1x Kinetics Buffer to ensure a consistent buffer composition for all concentrations in the series.

SA biosensors were loaded with biotinylated meGFP-tagged monobody for 120s to 600s to a density of 0.8 to 1.3 response units to ensure the loading density remained in the linear regime. Loading was stopped early if necessary. SA sensors were then dipped into a 1x kinetics buffer for 120s to establish a baseline. Association was monitored by dipping sensors into wells containing 3xFlag-tagged Omicron BA.1 RBD for 120s and dissociation was monitored by dipping wells into wells containing 1x kinetics buffer for at least 160s. Titration experiments were performed at 25°C while rotating at 1000 rpm. To measure *K*_d_, steady-state and global kinetic fits were performed using the manufacturer’s software (ForteBio Data Analysis 9.0) assuming a 2:1 heterogeneous ligand binding model.

### In cellulo K_d_ determination by yeast titrations

Monobody sequences were amplified from the print plate using primers appending 5’ and 3’ overhangs to pCTCON2. Reactions were PCR purified and assembled into a pre-digested (NheI/BamHI) pCTCON2 backbone via gibson assembly. Plasmids were miniprepped and sequence validated before being chemically transformed into EBY100 using a Frozen-EZ yeast transformation II kit (Zymo).

A single colony was selected per clone, passaged twice in SDCAA, and induced in SGCAA at a final seeding density of 1 x 10^7^ cells/mL. Expression was induced for 18-20 hours at 30°C in a shaking incubator. Using anti-c-myc FITC (1:100) and Omicron/BA.1 RBD Spy-AF647 as staining reagents, 3 x 10^6^ yeast were labeled per staining condition in a 30uL volume according to a published protocol^2^, taking care to ensure staining volume was modified when necessary to avoid the ligand titration regime. Following incubation, yeast cells were washed once in PBSA and kept on ice as pellets. Just before flowing, yeast cells were resuspended in PBSA and passed through a 35μm mesh filter. Yeast cells were visualized using a Sony SH800S FACS instrument with 70 μm sorting chip.

### Omicron BA.1 RBD SSM library screening

The Omicron BA.1 library^33^ was ordered from Addgene (#1000000187) and propagated as recommended. Maxiprepped plasmids were digested with ScaI prior to transformation into electrocompetent EBY100 to improve the efficiency of homologous recombination. The transformed library was passaged in SDCAA before being induced at 20°C for 36-48 hours. 2 x 10^7^ induced yeast cells per sort were washed three times with PBSA. For the screen against s14335, yeast cells were stained with anti-c-myc FITC (1:100) and 360nM of biotinylated, His-Avi-tagged s14335 monobody in a 100uL volume. For the screen against s19382, yeast cells were stained with anti-HA allophycocyanin (APC; 1:100; BioLegend) and 16nM sfGFP-tagged s19382 monobody in a 520uL volume. Cells were sorted as described previously. Following sorting, plasmids were miniprepped and prepared as before. Samples were submitted for either paired end 250 at Azenta (South Plainfield, NJ) or paired end150 at Neochromosome (Long Island City, NY) for NGS sequencing.

BBMerge^48^ (V39.01) was used to merge demultiplexed reads, Cutadapt^49^ (V2.6) was used to trim adapters, with a maximum error rate of 1/60. Custom python scripts were used to filter and map barcodes, as well as to calculate enrichment scores^20,35^ from naive and enriched and for performing downstream analyses. Barcodes containing any per-base quality scores <30 (for paired end 300 reads) or <40 (for paired end 150 reads) were not considered.

### Cryo-EM sample preparation

A 3.5-fold molar excess of sfGFP-tagged monobody s19382 was added to SARS-CoV-2 Omicron prefusion stabilized Spike protein (Sino Biological Cat. V40589-V08H26) and incubated for overnight at 4°C. The sample was concentrated and purified by size-exclusion chromatography on an AKTA Micro system using a Superose 6 Increase 3.2/300 column that had been pre-equilibrated in buffer containing 25 mM Tris pH 7.5, 150 mM NaCl. Fractions containing complex were pooled and concentrated to 1.8 mg/mL.

### Cryo-EM sample vitrification and data acquisition

Sample was vitrified by application of 3 µL onto a QuantiFoil Gold 1.2/1.3 300 mesh grid, which was blotted and plunge-frozen in liquid ethane using a Vitrobot Mk4 operated at 4°C and 100% humidity (wait time 10 s, blot time 4 s, blot force 0). The vitrified sample was then imaged on a Titan Krios G3i electron microscope operated at 300 keV and equipped with a BioQuantum energy filter and a K3 Summit direct electron detector camera. Images were recorded at a 105,000x nominal magnification corresponding to a pixel size of 0.84 Å/pix using a 20 eV energy slit. Each image stack contained 40 frames with the dose fractionation on the specimen set to 1.31 electrons per Å^2^ for a total dose of 42.5 *e^-^*/Å^2^. Images were collected with a nominal defocus range of -0.8 to -2.8 µm.

### Cryo-EM data processing

All data were processed using CryoSPARC^62^. Raw movies were motion corrected, and their contrast transfer function (CTF) estimated using patch motion correction and patch CTF estimation, respectively. Images were filtered to include only those with a detected fit resolution better than 5Å. Initial particle picks using CryoSPARC’s blob picker were subjected to iterative rounds of reference-free 2D classification, and a subset of particles from the most well-defined 2D classes were used to generate two ab initio models with no symmetry imposed. One class contained Spike monomers that were primarily in the closed state while the other contained Spike dimers with clear extra density at the dimer interface between the Spike RBDs. The full particle stack from all non-junk 2D classes was then subjected to iterative rounds of heterogeneous refinement using this dimeric ab initio model and several junk volumes as references. This cleaned particle stack was used to generate a second particle stack using cryoSPARC’s implementation of Topaz^63^. This stack was also subjected to iterative 2D classification and heterogeneous refinement, and both particle stacks were combined, removing duplicate particles. Non-uniform refinement was performed on the cleaned particle stack using the dimeric ab initio model as a reference. In this reconstruction, two RBDs from each spike protein were well resolved, with electron density corresponding to eight molecules of s19382 connecting the four RBDs. The third RBD from each spike protein was poorly resolved. In order to improve the resolution of the 4:8 RBD:s19382 dimer interface and understand the molecular nature of dimerization, a soft mask was generated around this region of the reconstruction. A second mask was then generated around the rest of the Spike dimer and particle subtraction was performed to remove the signal not corresponding to the interface of interest. Local refinement was then performed to yield a final 3.16 Å map of the dimer interface (determined by the gold standard Fourier shell correlation threshold of 0.143). Map sharpening for model building was performed using EMReady^64^ v2.0 using default parameters and without specifying an input structure or mask.

### Model building and structural analysis

An initial model was generated in UCSF ChimeraX^65^ by rigid body fitting copies of the Omicron RBD from PDB 7U0N^66^ and a model of s19382 into the density. As expected, four RBD and eight s19382 protomers could be fit into the density. This initial model was subjected to iterative rounds of manual rebuilding in Coot^67^ and refinement in Phenix^68^, followed by molecular dynamics-assisted manual refinement with ISOLDE^69^ in ChimeraX and a final round of refinement in Phenix.

