## Supplementary Information for "ADAPT-M: A workflow for rapid, quantitative *in vitro* measurements of enriched protein libraries"

**Supplementary Note 1:**

In FACS experiments, cells are sequentially labeled, first with RBD and then with SA, with a wash in between to remove free RBD. This kind of sequential labeling is not possible for STAMMPPING experiments because of the surface chemistry of the microfluidic device. Open NA sites on the device (if bBSA passivation was incomplete) could compete with immobilized monobody for binding to RBD, while accessible bBSA could result in false-positive binding signal from the fluorescently labeled avidin probe, confounding accurate $K_D$ determination. Thus, the target protein must be pre-incubated with the avidin probe prior to introduction onto the device.

We first sought to use an iFluor 650 monomeric streptavidin conjugate (mSA650) to prevent avidity effects. However, we observed significant non-specific binding to empty chambers, meGFP-containing chambers, and to surfaces outside of the button region. This suggested at least one of three issues: (1) incomplete binding of RBD to mSA650 during incubation, (2) dissociation of RBD from mSA650 over the course of the measurements, and/or (3) excess open biotin sites on the device surface. To address these issues we (1) extended incubation of RBD-mSA650 to 40 hours, well over five half-lives recommended for proper binding equilibration, (2) simultaneously performed experiments using a standard, tetrameric NA-Dylight 650 (NA650) probe, which offers slower dissociation rates than mSA, and (3) experimented with increased blocking times to minimize available biotin sites, respectively. Increasing the length of blocking to two hours resulted in the largest reduction of non-specific binding, and swapping to a NA650 probe further improved selective patterning of button surfaces. Even so, overnight NA blocking with buttons open was necessary to reduce non-specific binding to acceptable levels. While this reduced non-specific binding of the NA probe, leaving the buttons open while flowing NA overnight resulted in minor cross-contamination across chambers, as assessed from meGFP expression levels in empty chambers. To minimize the effects of cross-contamination, we implemented a very conservative expression threshold. As a result, measured affinities are primarily driven by the printed designs, not by any contaminating designs. Furthermore, since many replicates are printed across the device, each replicate surrounded by different neighboring sequences, any artifacts from small amounts of contamination average out across the replicates, reducing their impact on the overall measurements.

Given the non-specific binding to "empty" chambers, we applied a t-test to compare cy5 fluorescence intensities in aggregated chambers for each design against those of all empty chambers. Designs with $p > 0.05$ were classified as binding below the limit of detection, and apparent dissociation constants ($K_{d,\,app}$) were reported for all other designs. The dynamic range of measured $K_{d,app}$s was limited, spanning 82 nM to 1.1 µM, with poor resolution among binders in the ~100 nM range. All designs that showed detectable binding in this set of measurements were selected for validation with a monovalently labeled target (**Figure 2f**). Additionally, a subset of designs that appeared to bind below the limit of detection were included as negative controls, and to determine whether binding could be measured with an improved limit of detection (**Supplementary Figure S9**).

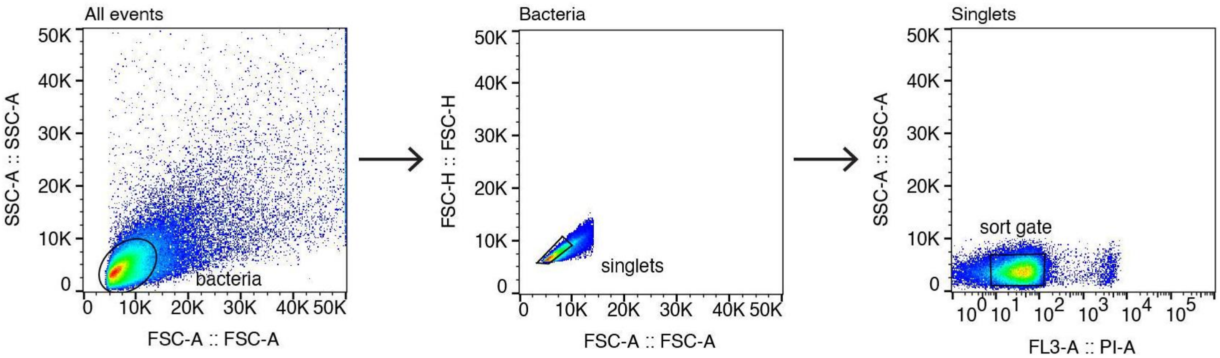

**Figure S1.** FACS plots showing sorting strategy for isolating single bacterial cells on a Sony SH800. Bacterial cells were stained with PI prior to sorting to distinguish live/dead cells. Left: Bacterial cells were first gated by SSC and FSC. Center: FSC-H vs FSC-A was used as a gate for singlets. Right: $PI^-$ cells were selected for sorting into individual wells of a multiwell plate and outgrowth. Approximate gates are outlined in black.

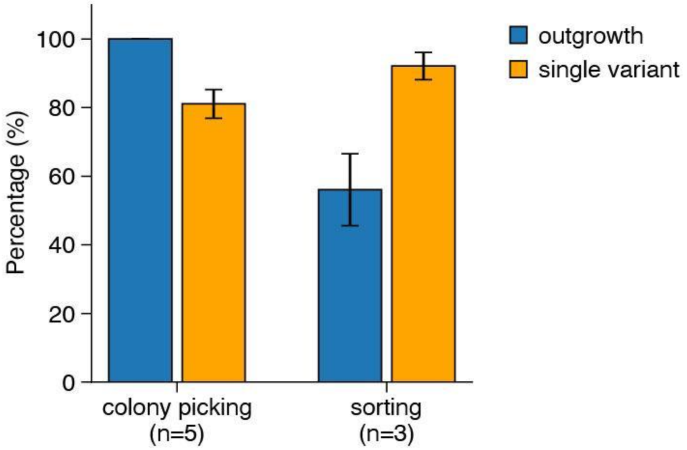


**Figure S2**. Bar plots showing cell outgrowth and single variant isolation rates for manual colony picking versus single cell sorting. Data represent mean values; error bars indicate standard error from biological replicates.

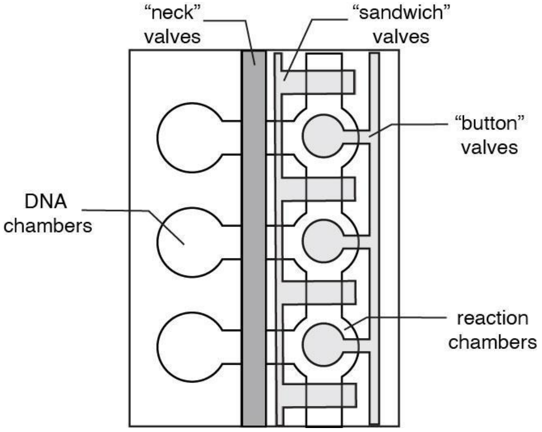


**Figure S3.** Cartoon schematic of chambers and valves on the microfluidic device. Putative binders are expressed in DNA chambers that are precisely aligned to linear expression templates during device bonding. These DNA chambers are separated from reaction chambers by "neck" valves, where putative binders are immobilized and binding is surveyed. "Sandwich" valves separate reaction chambers from one another to prevent cross-chamber contamination, and “button” valves permit both selective surface patterning of reaction chambers with anti-GFP nanobodies and protection of binding interactions during imaging.

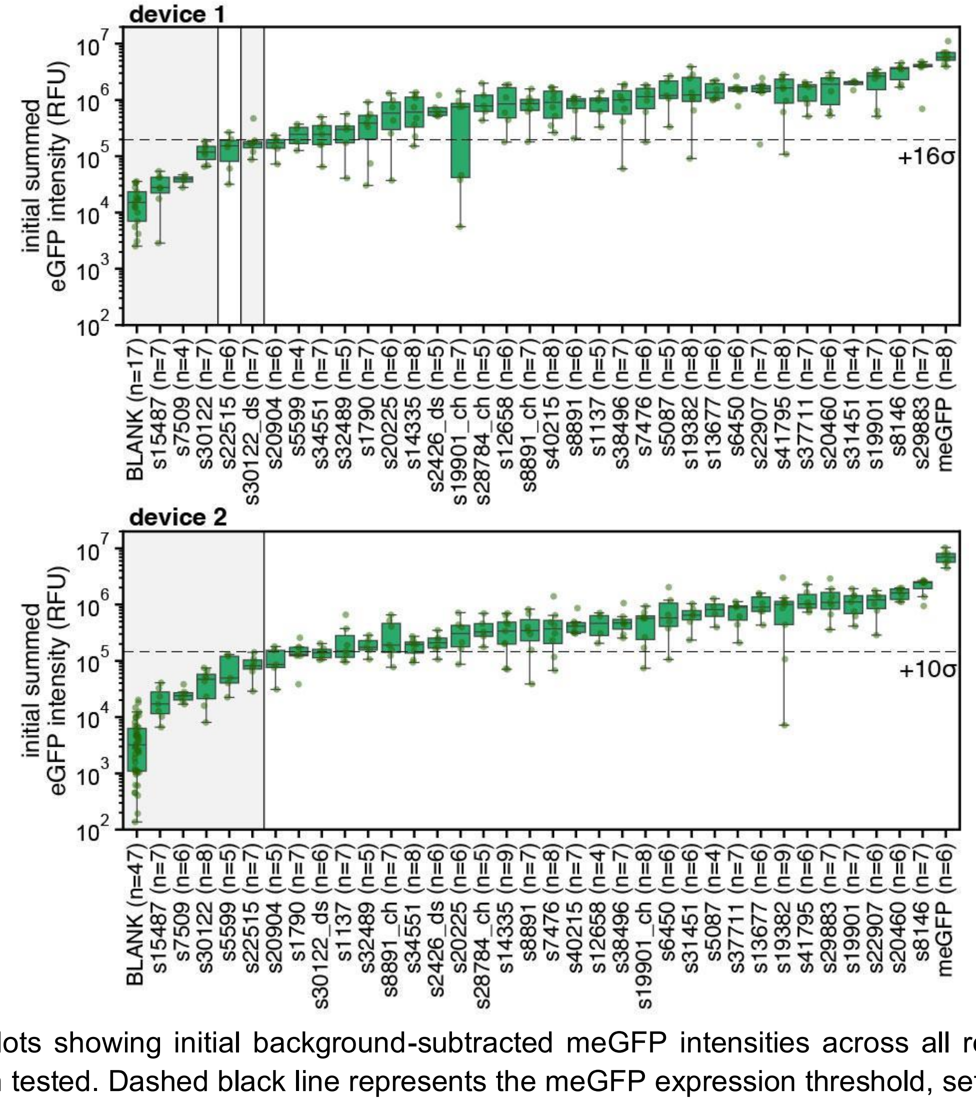


**Figure S4.** Boxplots showing initial background-subtracted meGFP intensities across all replicates for each monobody design tested. Dashed black line represents the meGFP expression threshold, set as the annotated number of standard deviations above the mean meGFP intensity across blank chambers for each device. Replicates with meGFP intensities below the threshold were not used for $K_d$ estimation. Grey shading indicates designs without at least 2 replicates expressing above this threshold.

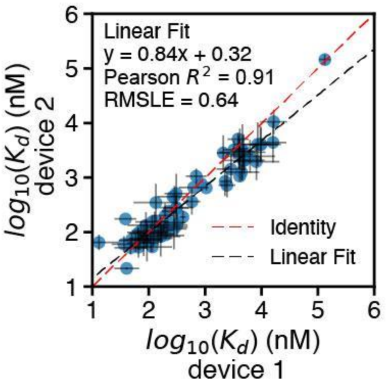


**Figure S5.** Pairwise comparison of per variant $K_d$s for all designs with binding above the limit of detection across 2 experiments. Points indicate median affinities (± SEM) for each design. Dashed red line indicates the identity line (1:1) and the dashed black line represents the best fit linear regression.

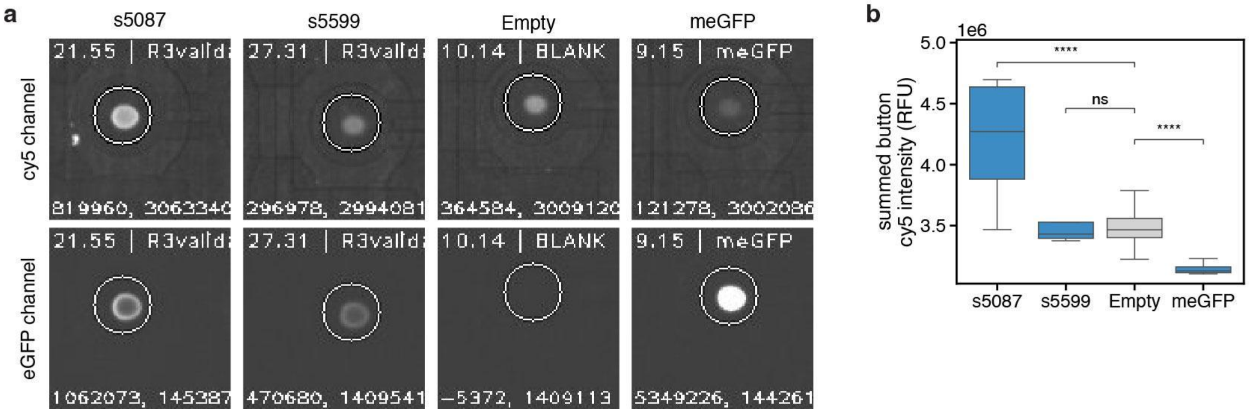


**Figure S6.** Defining experimental limit of detection. **a**. Images showing measured cy5 intensities (top) and meGFP intensities (bottom) for a weakly binding (s5087) monobody-containing chamber and monobody-containing chambers with binding below the limit of detection (s5599), along with corresponding intensities for meGFP-only and empty chambers. Contrast is uniformly enhanced for optimal visibility. Empty chambers were used to calculate the limit of detection for monobody-containing chambers, providing a strict binding threshold since meGFP-only containing chambers typically bind much more weakly than empty chambers. **b**. Distribution of summed button cy5 intensities at the highest RBD concentration for meGFP-containing and empty chambers along with weakly binding (s5087) and nonbinding (s5599) monobody-containing chambers. A two-sided t-test was applied, **** denotes $p < 0.0001$, ns denotes $p > 0.05$.

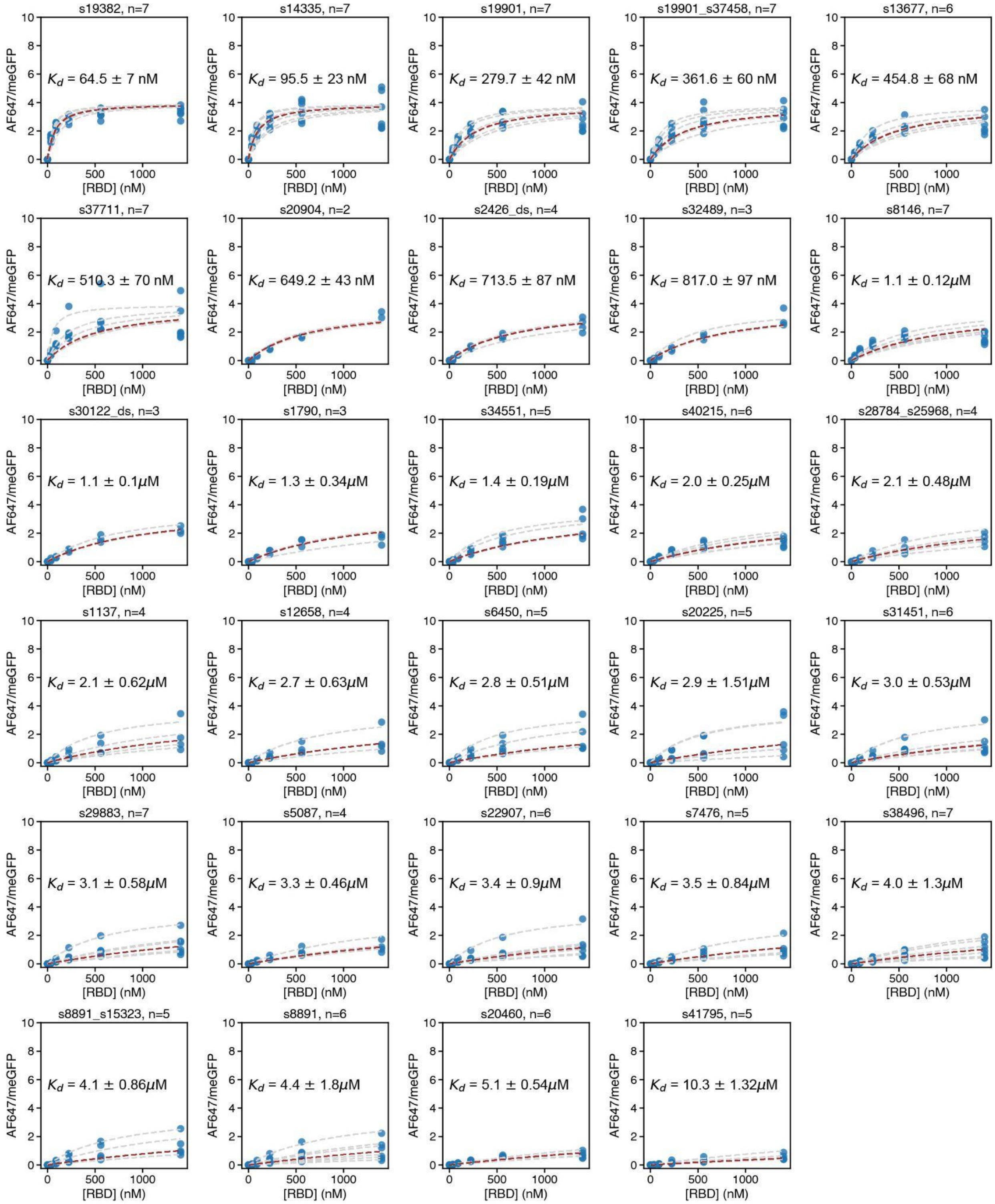


**Figure S7**. Concentration-dependent binding curves for monobody library measured against monovalent, SpyTag/Catcher-AF647-labeled Omicron/BA.1 RBD. Light grey dashed lines indicate per-chamber Langmuir isotherm fits. The median $K_d$ is annotated, and the dashed red line indicates the fit returning the median $K_d$. Blue markers indicate measured intensity ratios at each RBD concentration.

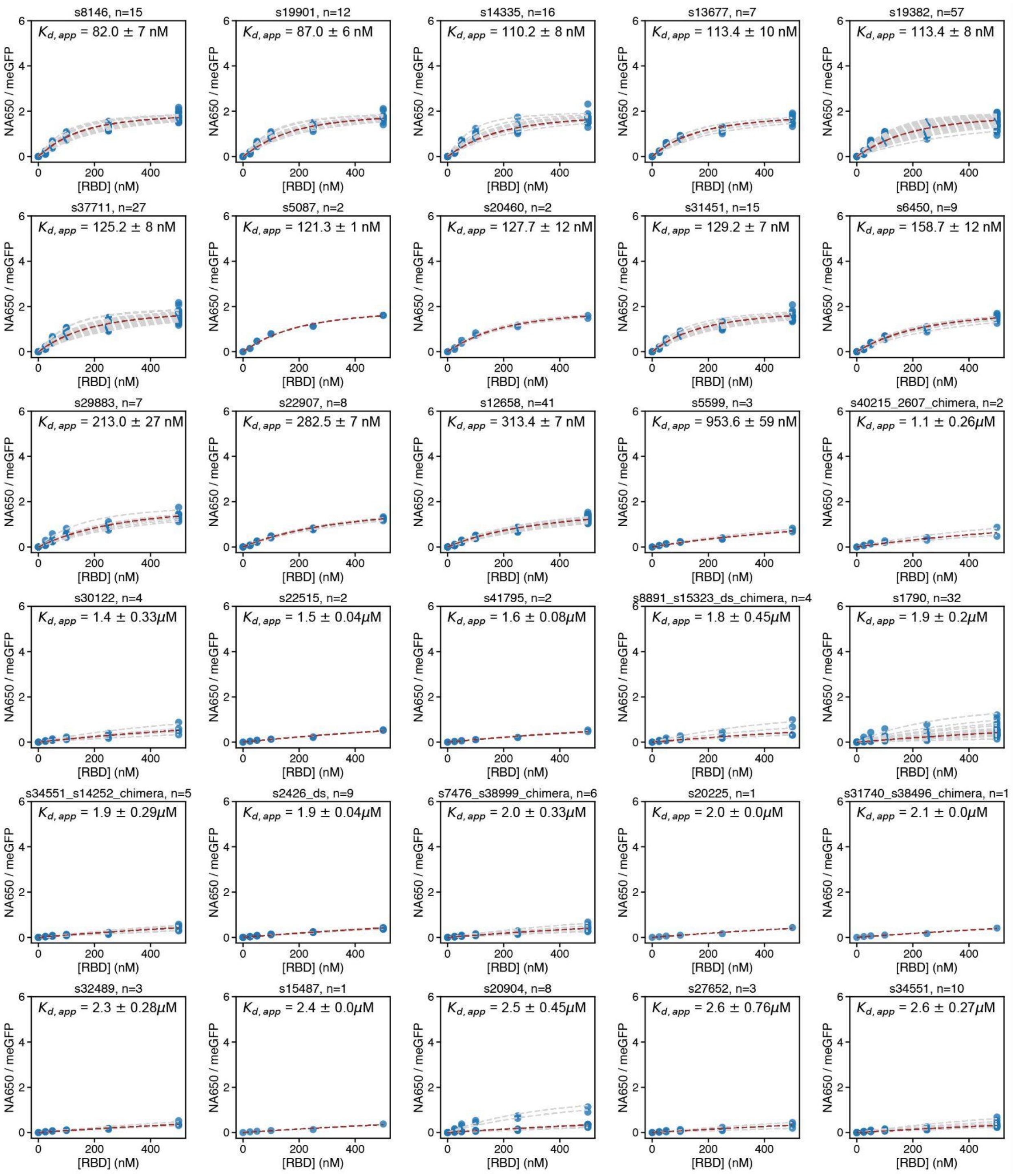

s8146, n=15
$K_{d,\,app}$ = 82.0 ± 7 nM
s19901, n=12
$K_{d,\,app}$ = 87.0 ± 6 nM
s14335, n=16
$K_{d,\,app}$ = 110.2 ± 8 nM
s13677, n=7
$K_{d,\,app}$ = 113.4 ± 10 nM
s19382, n=57
$K_{d,\,app}$ = 113.4 ± 8 nM
s37711, n=27
$K_{d,\,app}$ = 125.2 ± 8 nM
s5087, n=2
$K_{d,\,app}$ = 121.3 ± 1 nM
s20460, n=2
$K_{d,\,app}$ = 127.7 ± 12 nM
s31451, n=15
$K_{d,\,app}$ = 129.2 ± 7 nM
s6450, n=9
$K_{d,\,app}$ = 158.7 ± 12 nM
s29883, n=7
$K_{d,\,app}$ = 213.0 ± 27 nM
s22907, n=8
$K_{d,\,app}$ = 282.5 ± 7 nM
s12658, n=41
$K_{d,\,app}$ = 313.4 ± 7 nM
s5599, n=3
$K_{d,\,app}$ = 953.6 ± 59 nM
s40215_2607_chimera, n=2
$K_{d,\,app}$ = 1.1 ± 0.26$\mu$M
s30122, n=4
$K_{d,\,app}$ = 1.4 ± 0.33$\mu$M
s22515, n=2
$K_{d,\,app}$ = 1.5 ± 0.04$\mu$M
s41795, n=2
$K_{d,\,app}$ = 1.6 ± 0.08$\mu$M
s8891_s15323_ds_chimera, n=4
$K_{d,\,app}$ = 1.8 ± 0.45$\mu$M
s1790, n=32
$K_{d,\,app}$ = 1.9 ± 0.2$\mu$M
s34551_s14252_chimera, n=5
$K_{d,\,app}$ = 1.9 ± 0.29$\mu$M
s2426_ds, n=9
$K_{d,\,app}$ = 1.9 ± 0.04$\mu$M
s7476_s38999_chimera, n=6
$K_{d,\,app}$ = 2.0 ± 0.33$\mu$M
s20225, n=1
$K_{d,\,app}$ = 2.0 ± 0.0$\mu$M
s31740_s38496_chimera, n=1
$K_{d,\,app}$ = 2.1 ± 0.0$\mu$M
s32489, n=3
$K_{d,\,app}$ = 2.3 ± 0.28$\mu$M
s15487, n=1
$K_{d,\,app}$ = 2.4 ± 0.0$\mu$M
s20904, n=8
$K_{d,\,app}$ = 2.5 ± 0.45$\mu$M
s27652, n=3
$K_{d,\,app}$ = 2.6 ± 0.76$\mu$M
s34551, n=10
$K_{d,\,app}$ = 2.6 ± 0.27$\mu$M
NA650 / meGFP
[RBD] (nM)

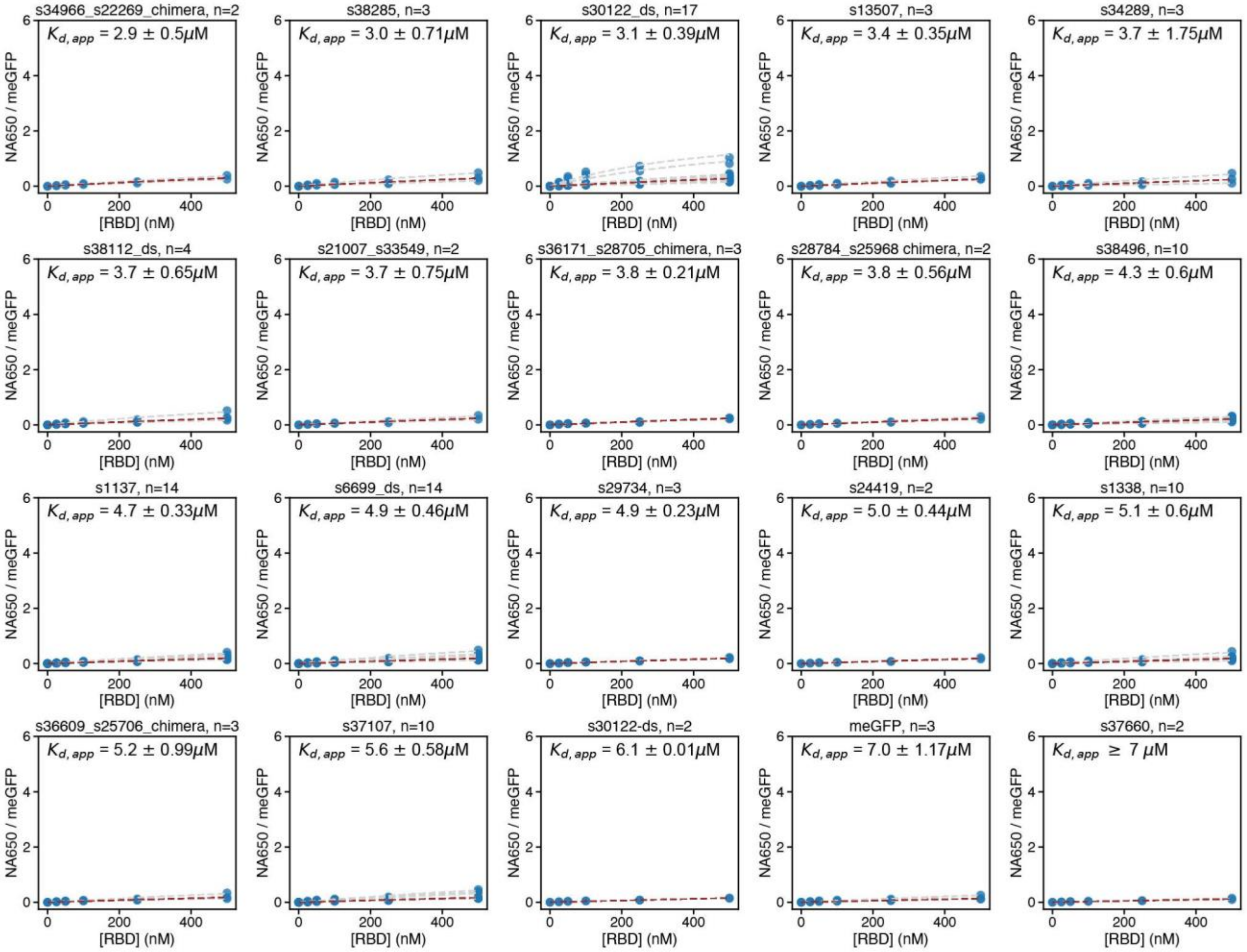


**Figure S8.** Concentration-dependent binding curves for monobody library measured against a tetrameric, neutravidin-iFluor650-labeled RBD. Light grey dashed lines indicate per-chamber Langmuir isotherm fits. The median $K_{d,app}$ is annotated, and the dashed red line indicates the fit returning this $K_{d,app}$. Blue markers indicate measured intensity ratios at each RBD concentration.

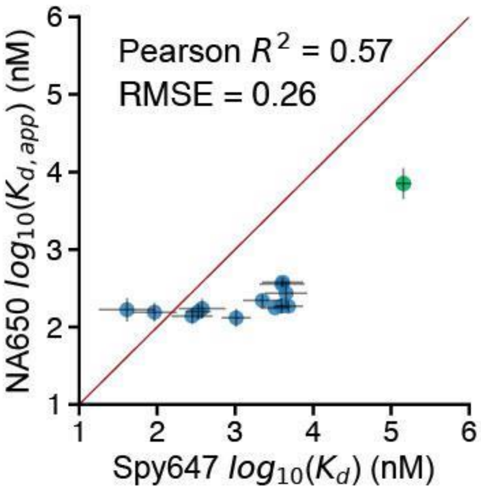


**Figure S9.** $K_d$ values of avid neutravidin-based STAMMPPING measurements are compressed compared to measurements made with monovalent RBD.

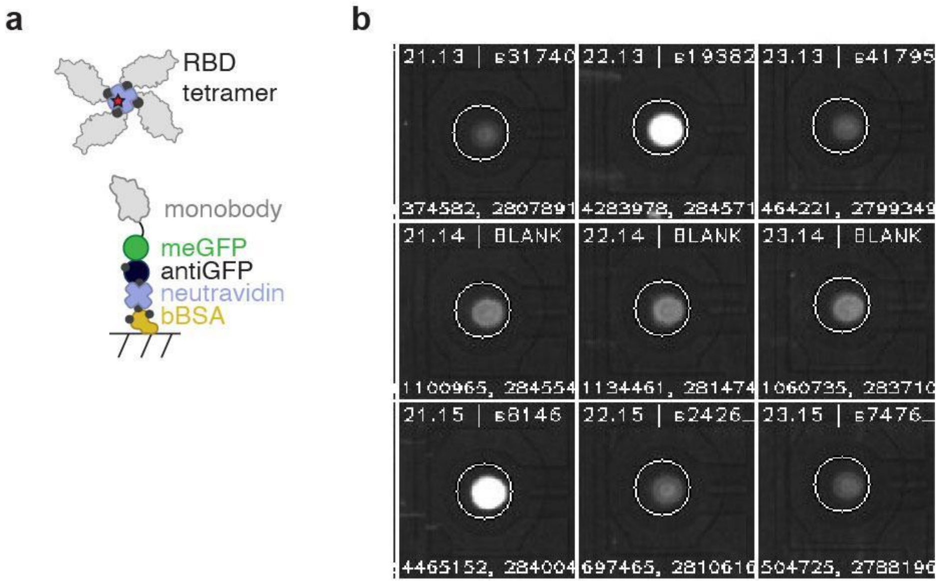


**Figure S10**. Images showing cy5 intensities for subset of chambers across microfluidic device using neutravidin-647 RBD target.

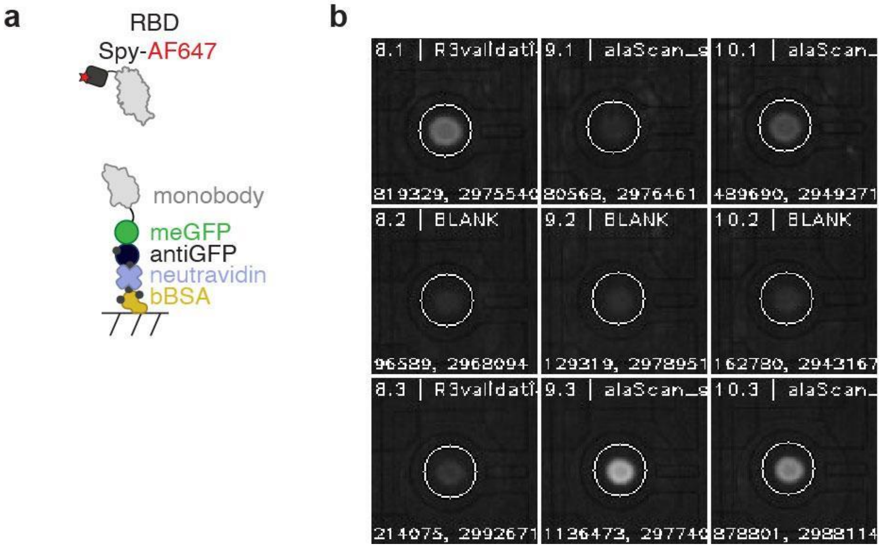


**Figure S11**. Images showing cy5 intensities for subset of chambers across microfluidic device using AF-647 SpyTag RBD target.

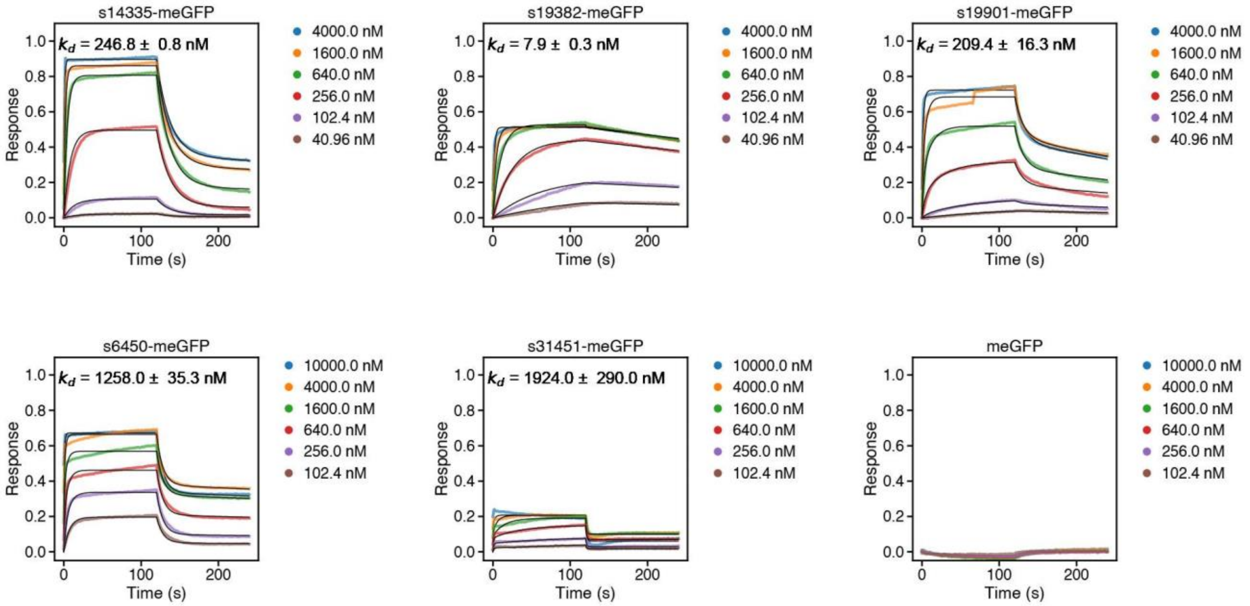


**Figure S12.** Kinetic binding traces and global kinetic fits measured by BLI.

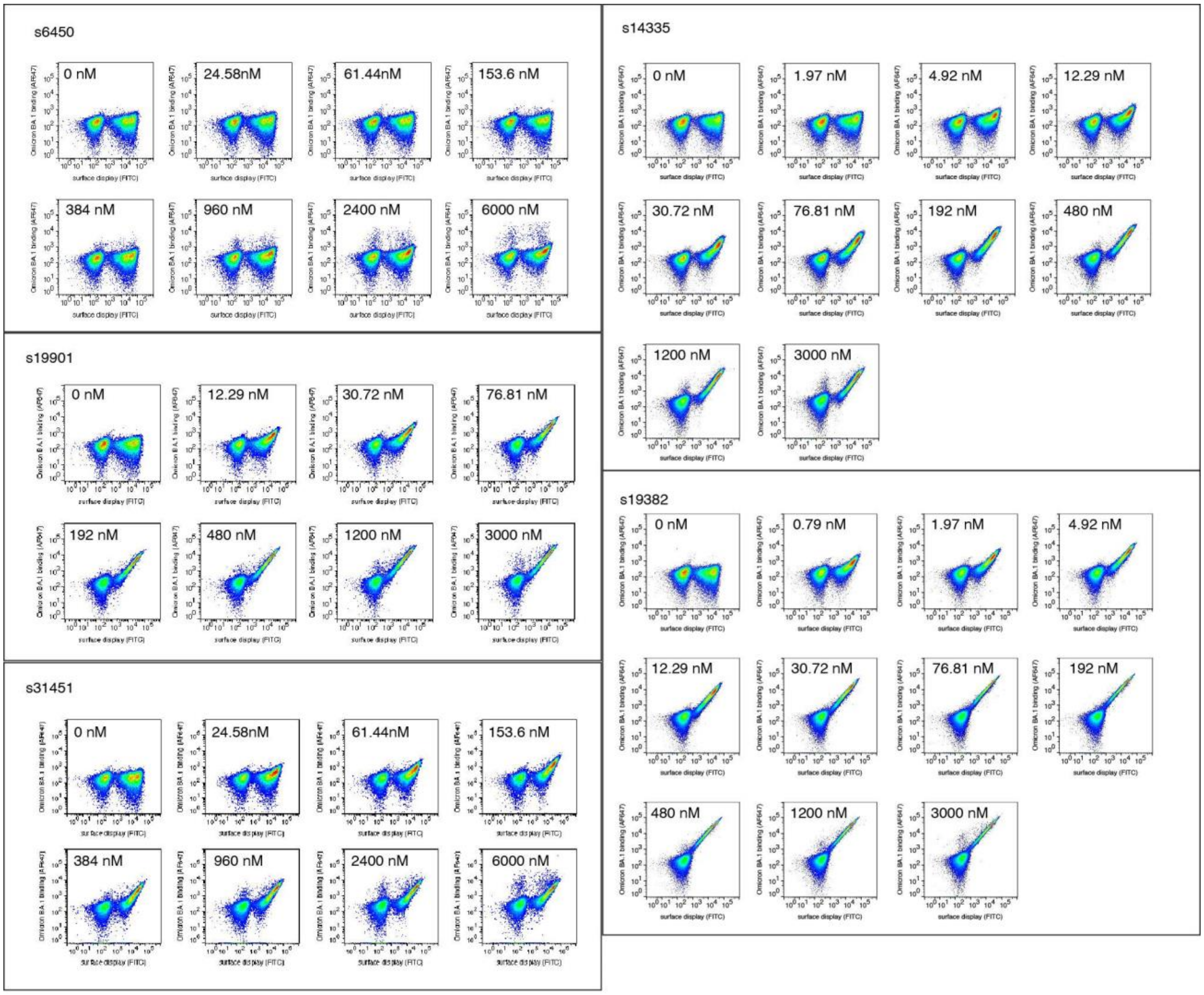


**Figure S13.** FACS plots from one replicate of titration series for 5 designs. Staining concentrations are annotated.

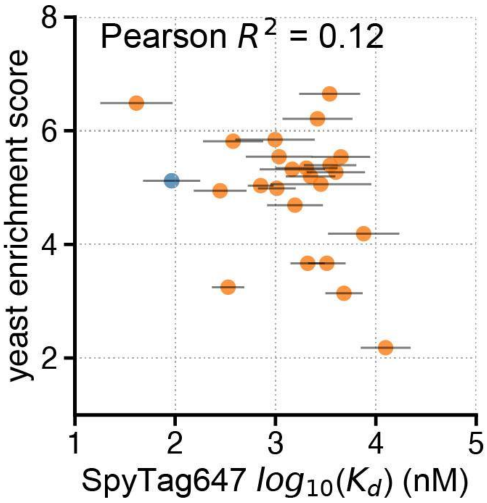


**Figure S14.** Log enrichment scores from yeast surface display plotted as a function of $K_d$ measured via STAMMPPING. Individual designs represented as orange scatter points; s14335 shown in blue.

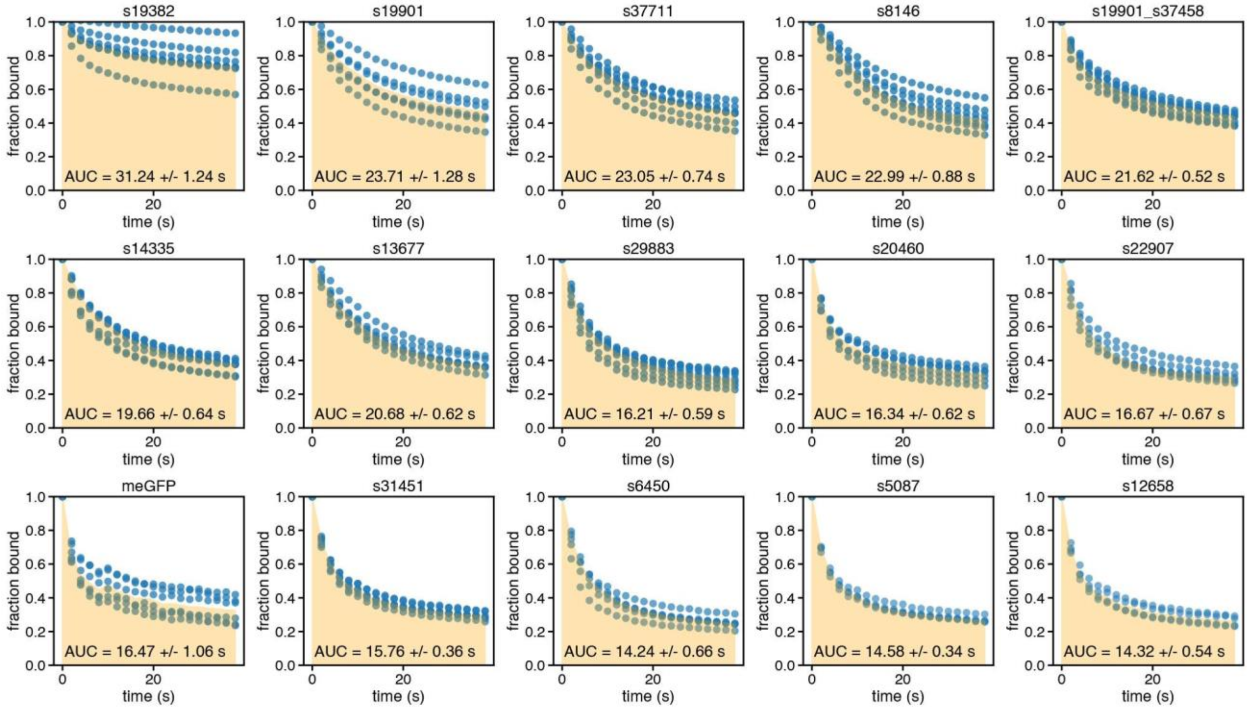


**Figure S15.** Time-dependent dissociation curves for monobody library measured against Omicron RBD via STAMMPPING. Blue markers indicate normalized, measured AF647 intensities at each time point. AUC for median dissociation at each time point is shaded in orange for each designed binder. Median AUCs (± SEM) are annotated.

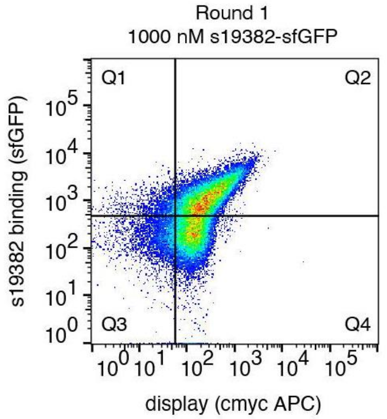


**Figure S16.** Representative flow cytometry dot plots of Omicron BA.1 SSM yeast surface display screening against s19382. sfGFP-tagged s19382 was used as a primary stain, and anti-HA APC was used to detect the displaying population. Target staining concentrations are indicated. Q2 represents the approximate sort gate.

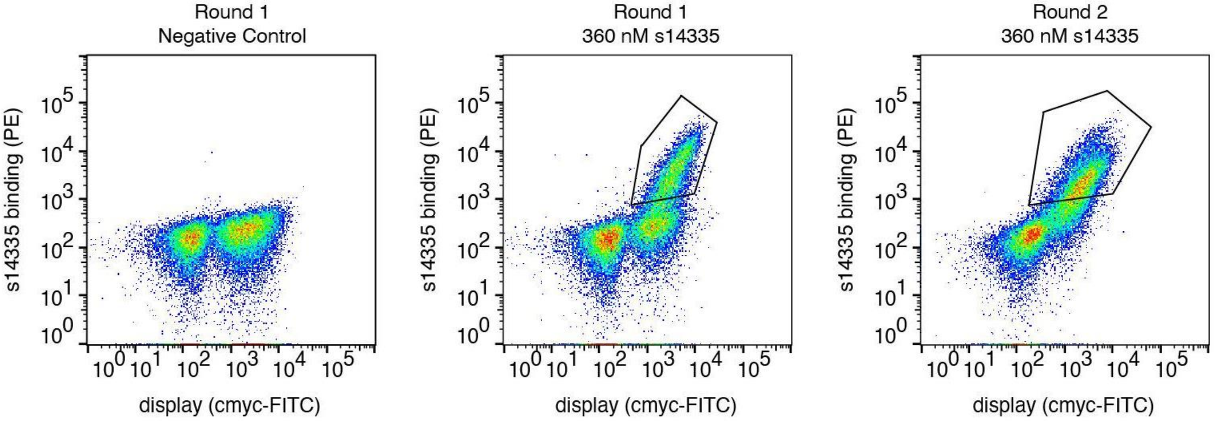


**Figure S17.** Representative flow cytometry dot plots of Omicron BA.1 SSM yeast surface display screening against s14335. Yeast labeled with secondary stains and without any putative binder were used as a negative control to set binding gates (left). His- Avi-tagged s14335 was biotinylated and used as a primary stain, streptavidin-PE served as a secondary for detection of s14335-bound cells. Anti-cmyc FITC was used to detect the displaying population. Target staining concentrations are indicated. Approximate sort gates for 2 iterative rounds of enrichment (center, right) are outlined in black.

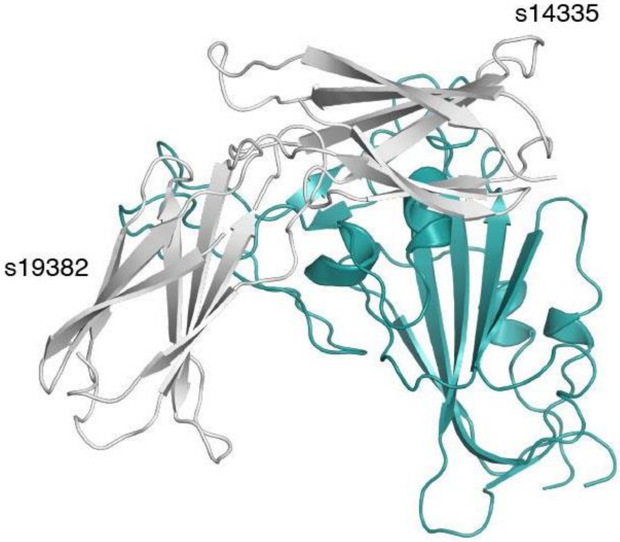


**Figure S18.** s14335 and s19382 design models (grey, labeled) binding to Omicron BA.1 RBD (teal).

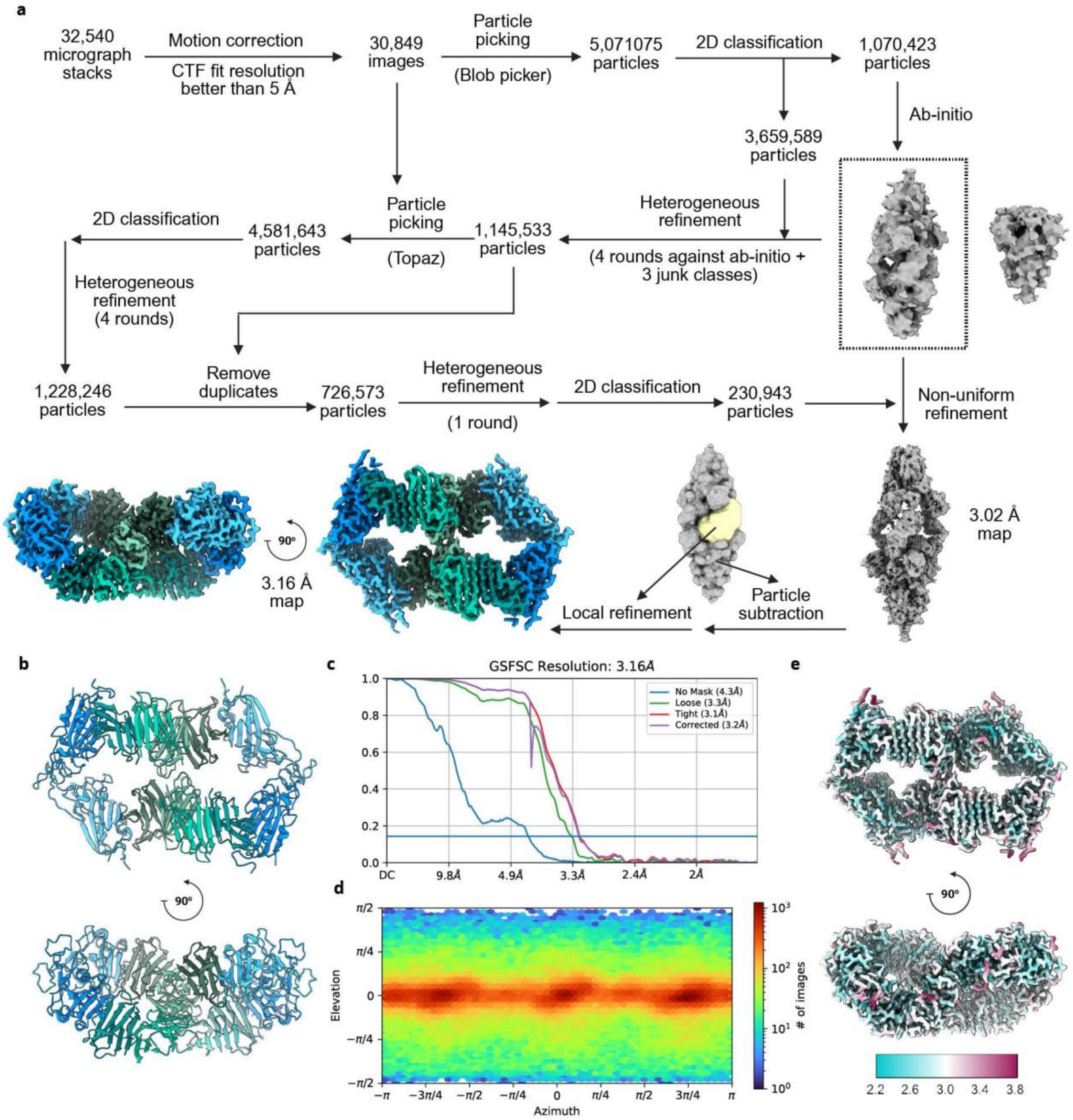


**Figure S19.** Cryo-EM workflow for structure determination of the s19382-RBD (Omicron) complex. **a** Schematic of the cryo-EM data processing workflow for the s19382-Omicron Spike sample. RBDs in the final reconstruction are colored in shades of blue and s19382 protomers are colored in shades of green. **b** Cartoon representation of the cryo-EM structure of the s19382-RBD (Omicron) complex, colored as in a. **c** Global resolution estimate based off the Fourier Shell Correlation (FSC) between two half datasets. **d** Heat map showing the overall distribution of assigned particle orientations in the final reconstruction. **e** Final reconstruction colored by local resolution.

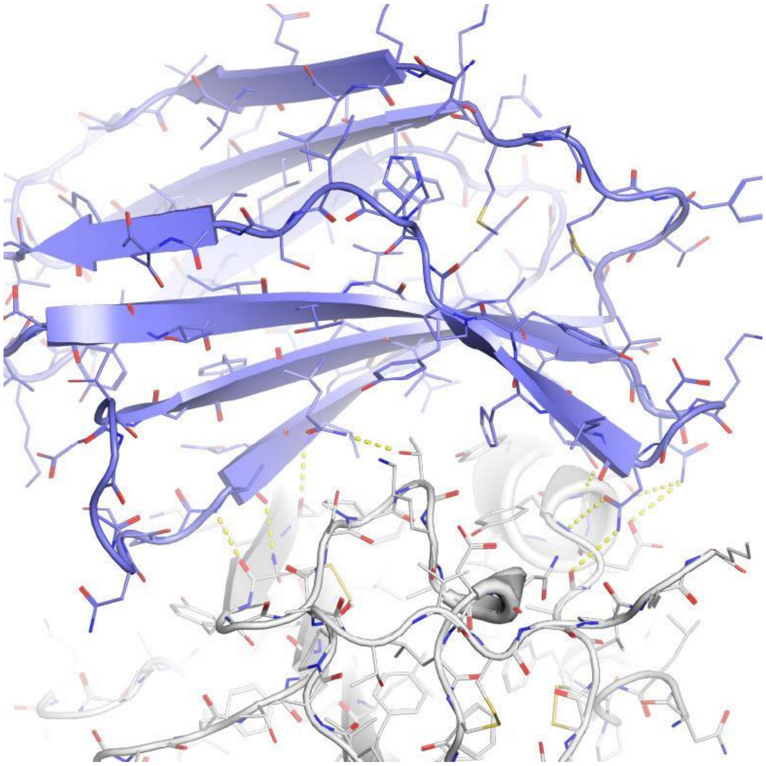

**Figure S20.** Close-up view of the binding interface of the s19382-RBD complex. RBD is shown in grey and s19382 is shown in purple. Yellow dashes denote polar contacts.

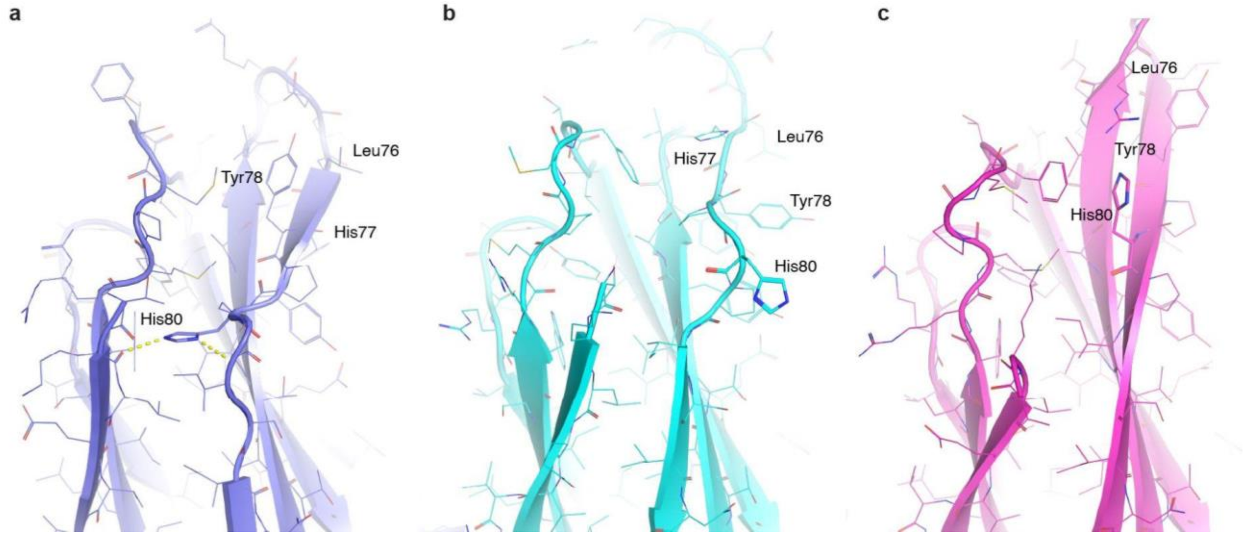


**Figure S21.** His80 side chain packing in experimental, designed, and predicted s19382 models. **a** Cryo-EM structure of s19382, showing His80 contributes to packing between βA and βB. **b** Design model for s19382. His80 is solvent-facing, resulting in a βG register shift as compared to the experimental model. **c** AlphaFold3 prediction of s19382. His80 is more solvent-exposed than in the experimental model and does not contribute to packing of βA and βG.

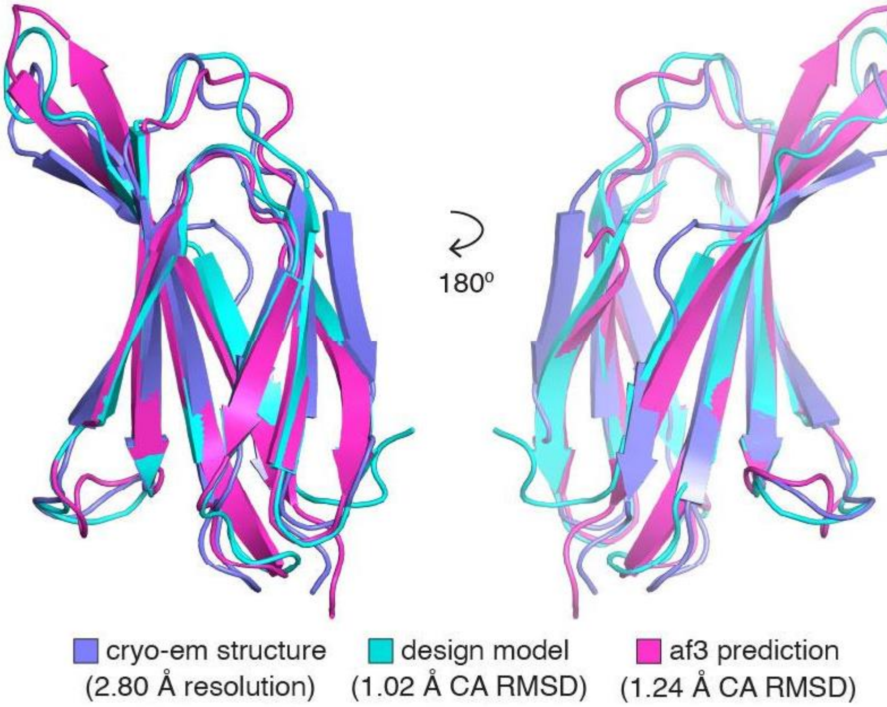


**Figure S22.** Overlay of experimental, predicted, and designed structures of monobody s19382. Cryo-EM model of s19382 is shown in purple, design model is shown in cyan, and AlphaFold3 prediction is shown in magenta.

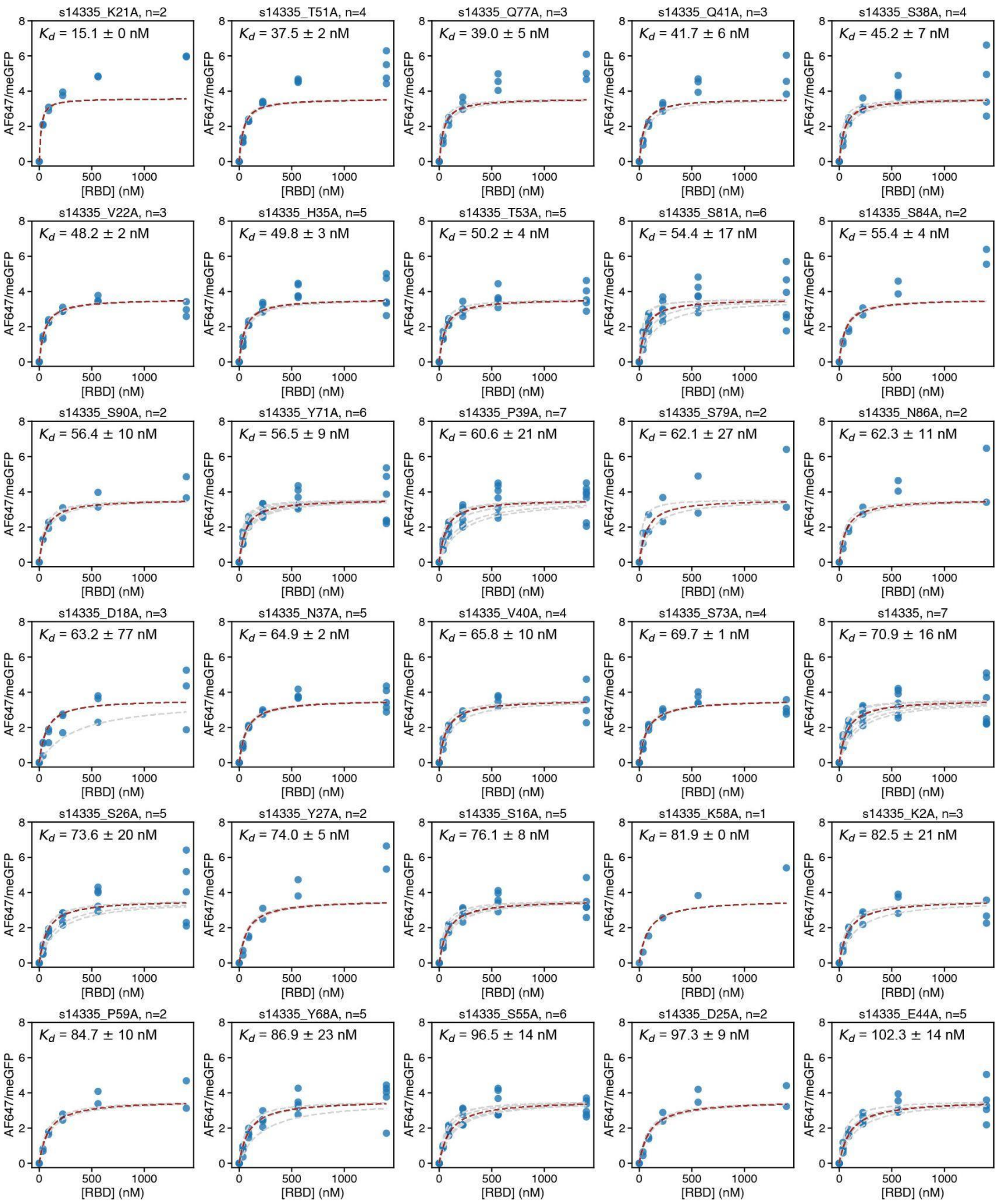
s14335_K21A, n=2
$K_d$ = 15.1 ± 0 nM
s14335_T51A, n=4
$K_d$ = 37.5 ± 2 nM
s14335_Q77A, n=3
$K_d$ = 39.0 ± 5 nM
s14335_Q41A, n=3
$K_d$ = 41.7 ± 6 nM
s14335_S38A, n=4
$K_d$ = 45.2 ± 7 nM
s14335_V22A, n=3
$K_d$ = 48.2 ± 2 nM
s14335_H35A, n=5
$K_d$ = 49.8 ± 3 nM
s14335_T53A, n=5
$K_d$ = 50.2 ± 4 nM
s14335_S81A, n=6
$K_d$ = 54.4 ± 17 nM
s14335_S84A, n=2
$K_d$ = 55.4 ± 4 nM
s14335_S90A, n=2
$K_d$ = 56.4 ± 10 nM
s14335_Y71A, n=6
$K_d$ = 56.5 ± 9 nM
s14335_P39A, n=7
$K_d$ = 60.6 ± 21 nM
s14335_S79A, n=2
$K_d$ = 62.1 ± 27 nM
s14335_N86A, n=2
$K_d$ = 62.3 ± 11 nM
s14335_D18A, n=3
$K_d$ = 63.2 ± 77 nM
s14335_N37A, n=5
$K_d$ = 64.9 ± 2 nM
s14335_V40A, n=4
$K_d$ = 65.8 ± 10 nM
s14335_S73A, n=4
$K_d$ = 69.7 ± 1 nM
s14335, n=7
$K_d$ = 70.9 ± 16 nM
s14335_S26A, n=5
$K_d$ = 73.6 ± 20 nM
s14335_Y27A, n=2
$K_d$ = 74.0 ± 5 nM
s14335_S16A, n=5
$K_d$ = 76.1 ± 8 nM
s14335_K58A, n=1
$K_d$ = 81.9 ± 0 nM
s14335_K2A, n=3
$K_d$ = 82.5 ± 21 nM
s14335_P59A, n=2
$K_d$ = 84.7 ± 10 nM
s14335_Y68A, n=5
$K_d$ = 86.9 ± 23 nM
s14335_S55A, n=6
$K_d$ = 96.5 ± 14 nM
s14335_D25A, n=2
$K_d$ = 97.3 ± 9 nM
s14335_E44A, n=5
$K_d$ = 102.3 ± 14 nM
AF647/meGFP
[RBD] (nM)

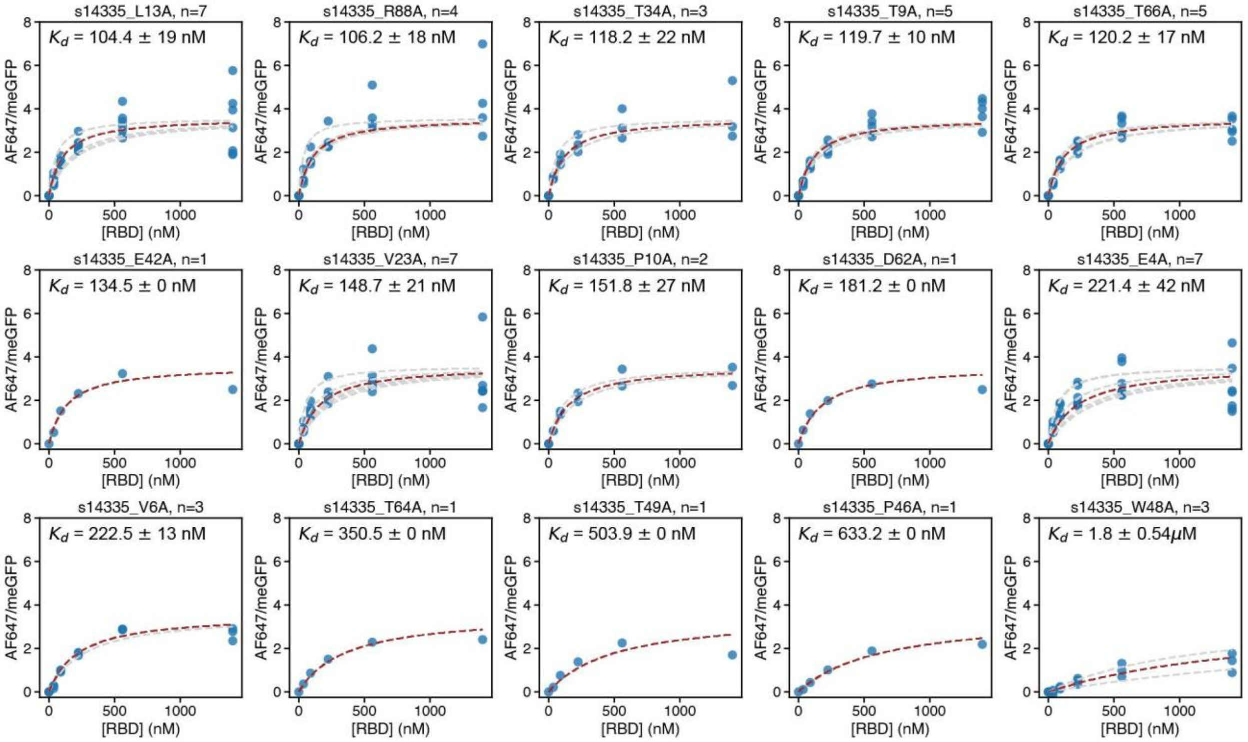


**Figure S23.** Concentration-dependent binding curves for s14335 alanine variants. Light grey dashed lines indicate per-chamber Langmuir isotherm fits. The median $K_\mathrm{d}$ is annotated, and the solid red line indicates the fit returning this $K_\mathrm{d}$. Blue markers indicate measured intensity ratios at each RBD concentration.

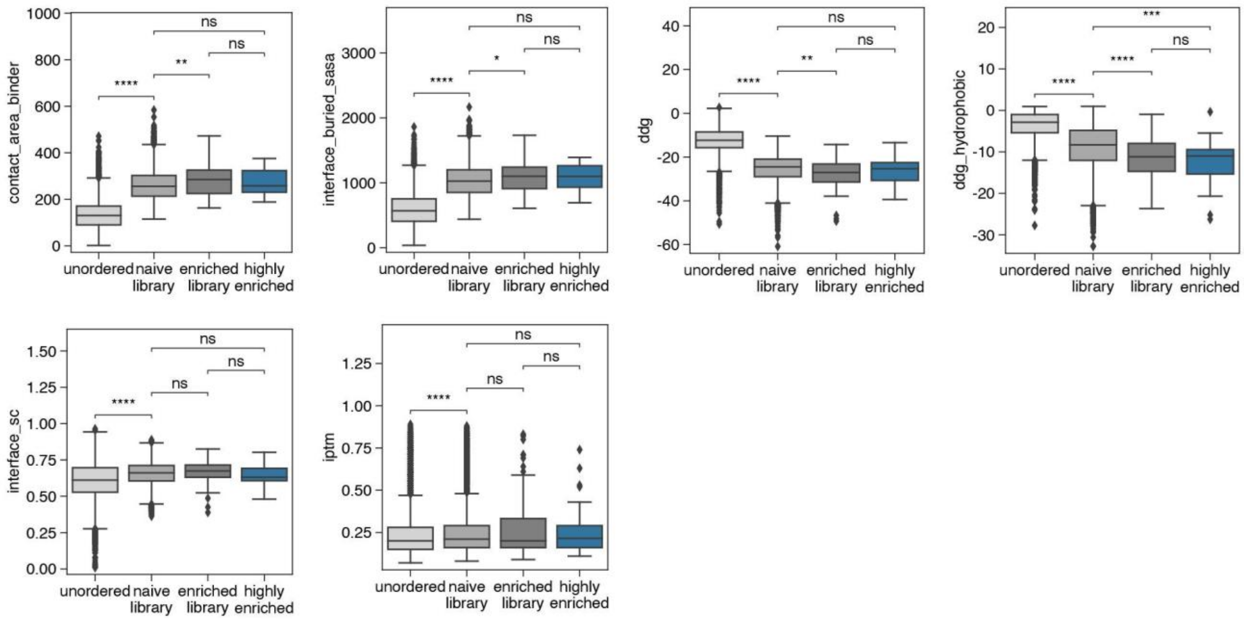

**Figure S24.** Spread of scores for a range of computational metrics compared across the ordered, enriched, and highly enriched library subsets. Two-sided t-tests were performed; ** denotes p<0.01, *** denotes p<0.001; **** denotes p<0.0001.

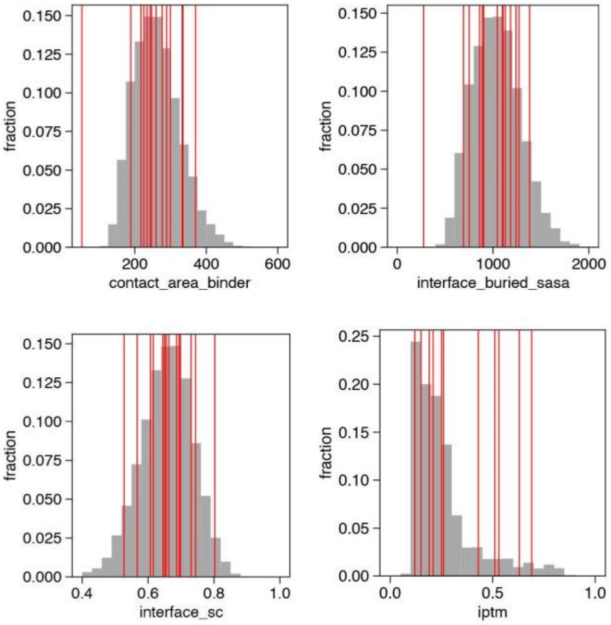


**Figure S25.** Histogram demonstrating spread of computational metrics (Rosetta and alphafold2) across the designed library (grey). Red vertical lines denote STAMMPPING validated binders.

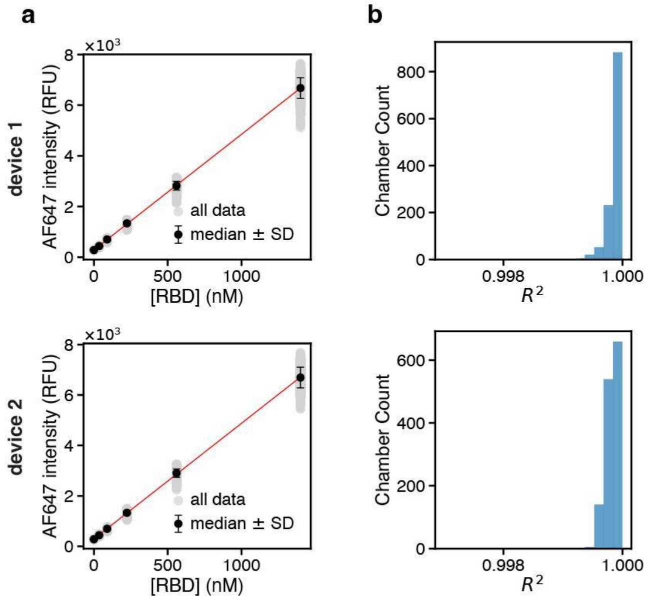

**Figure S26.** Calibration curves relating AlexaFluor-647 fluorescence intensities to RBD concentration. **a.** Per chamber calibration curves across sample chambers used for analysis. Light grey markers denote median per chamber intensities following 500ms exposure. Black markers denote median intensity across all chambers, and error bars denote standard deviation. Red line represents the best fit linear regression. **b.** Goodness of fit distributions for calibration curves.

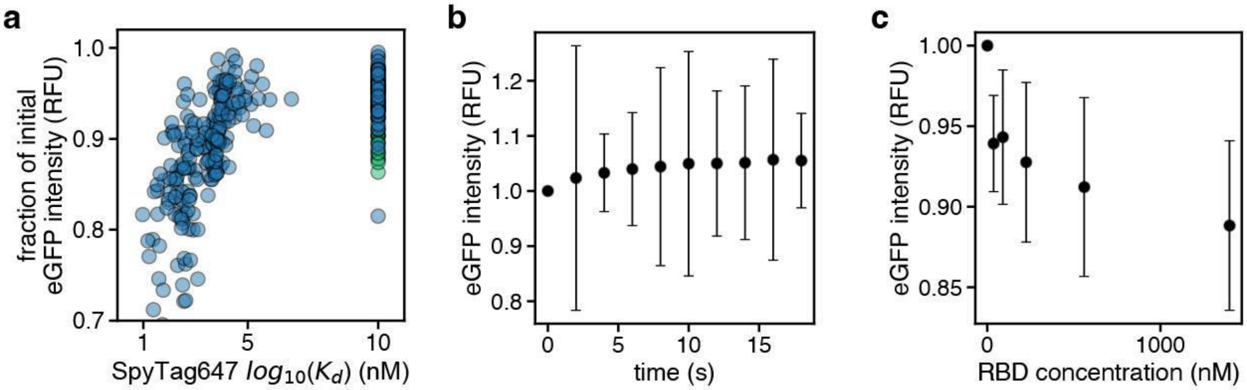


**Figure S27.** meGFP quenching and recovery over the course of binding and kinetics measurements. **a.** Scatter plot of fraction of initial meGFP fluorescence intensity as a function of per chamber $K_d$ measurements. Blue circles indicate monobody chambers, green circles indicate meGFP chambers. **b, c.** Black markers denote median intensity across all non-blank, non-culled sample chambers, and error bars denote standard deviation. **b.** Per chamber eGFP fluorescence as a function of RBD concentration during binding measurements. **c.** Per chamber eGFP fluorescence as a function of time during dissociation kinetics measurements.

**Table S1**: All printed sequences

| monobody ID | protein sequence | # colonies picked |
|---|---|---|
| s8146 | TKLEVVAATPTSLLIYWDHPMWDVDFYRITYGEVNGNSPVQSFNVPGS STVAWIYGLKPGVDYTITVYAYSGGSSQYFPSPISINYRTYNS | 6 |
| s19382 | TKLEVVAATPTSLLISWRMPMFTVDFYVIQYGETGGNSPVQVQVVPGST RTATISGLKPGVDYTITVYAYVQRDNLHYPHPISINYRTGSK | 23 |
| s41795 | TKLEVVAATPTSLLISWRHPMVYVDFYVITYGETHGNSPMQKFYVPGWK NTARISGLKPDVDYTITVYAYYSAPEWQKPSPISINYRTSGS | 1 |
| s22515 | TKLEVVAATPTSLLISWHFPAVTVDFYVITYGETGGNSPTDVFIVPGSAS TATISGLYDFVDYTITVYAFVESGSKGAAKPISINMTGGGE | 13 |
| s6450 | TKLEVVAATPTSLLISWDIPELQFDFYVITYGETFGNSPVQEFTVPGLATH ATISGLKPGVDYTITVYAYMSGSSYGSMIPISINYRTYGE | 4 |
| s5599 | TKLEVVAATPTSLLISWDYPWYIFDFYRITYGEVGGNMPMQEYEVPGQI | 2 |

| | | |
|---|---|---|
| | STATISGLKPGVDYTITVYAYVSIVEYYAPSPISINYRTRSG | |
| s40215_2607 | TKLEVVAATPTSLLISWDFPAVTIIFYFIEYGSTHGRDPIQSFRVPGSKST ATISGLKPGVDYTITVYAYHQRDNDHTPSPISINYRTGSG | 1 |
| s8891_s15323_ ds | TKLEVVAATPTSLLISWDAPAVWVFFYRITYGETGGNSPVQESFVPGDM STATISGLKPGVLYLITVYTMYHKNDEYPPNYHSPIFITYRT | 3 |
| s12658 | TKLEVVAATPTSLLISWDAPAITVDWYTITYGETGGNMPMQSFFVPGSM STATISGLKRDVDYTITVYAQQITFEQYVPSPISINYRVGGE | 15 |
| s34551 | TKLEVVAATPTSLLISWDHPAVDVNFYHITYGETGGNSPTYSFFVPGSV MTATISGLKPGVDYTITVYAHAASRYQAFPSPISINYRTGSP | 7 |
| s30122 | TKLEVVAATPTSLLISWDFPAVTFDFYVITYGEVGGNSPVQEFVVPGSVS TATISGLKPYVDYTITVYAWYYRSDGGWGSPSSMKGHGSPG | 3 |
| s19901 | TKLEVVAATPTSLLISWHWPAQTVDFYVITYGETGGNSPMQKFRVPGW VNTATISGLKPDVDYTITVYAHVVLPGHYSPPPISINYRPGPG | 6 |
| s7476_s38999 | TKLEVVAATPTSLLISWDAPAVDFVYYVITYGETHGNSPVQEFRVPGRLS HATISGLKPGVDYTITVYAFYHSDDSYPAEFMHPISINYRT | 2 |
| s29883 | TKLEVVAATPTSLLISWDMPMVTVVFYKITYGETGGNSPMQEFFVPGFL NTAKISGLKPGVDYTITVYAWLTGDEYYYSSPISINYRTESE | 2 |
| s1790 | TKLEVVAATPTSLLISWDMPMVWVDFYVITYGEQGGNSPKWKFIVPGST STATISGLKPGVDYTITVYAYAYVDHSVKPIMWNPRFQSFGS | 17 |
| s37711 | TKLEVVAATPTSLLISWDASQNHEYDFYRITYGQIGGVFPQYRFTVPGW TNTATITGLKPGVDYTITVYAYHKDGYLGNAFSSFNYRVGGG | 11 |
| s13677 | TKLEVVAATPTSLLISWDAERNWVLFYIITYGETHGNKPVQSFRVPGWIS TAWITGLKPGVDYTITVYAAGSQDDGGAIWSPISINYRTGG | 2 |
| s32489 | TKLEVVAATPTSLLISWHVPQSFVLFYVITYGETGGNSPVYHFFVPGSKN TATISGLKPGVDYTITVYAWQGGKDHFYGDSSGSYEGRPSS | 2 |
| s20904 | TKLEVVAATPTSLLISWDAEAYTVDFYLITYGETGGNSPMQEFIVPGRVT TATISGLKPGVDYTITVYAFYYGHGEGSWSQSGGWGYYNEG | 4 |
| s2426_ds | TKLEVVAATPTSLLISWDAPMVTVDFYFIVYGEVGGTSPLQWFTVPGSK NTATISGLKPGVDYTICIYAFRTGSDWFIGSGHCSDGKGGNT | 6 |
| s30122_ds | TKLEVVAATPTSLLISWDFPAVTFDFYVITYGEVGGNSPVQEFVVPGSVS TATISGLKPYVDYTICVYAWYYRSDGGWGSPSCMKGHGSPG | 8 |
| s34551_s14252 | TKLEVVAATPTSLLISWDHPAVDVNFYHITYGETGGNSPTYSFFVPGSV MTATISGLVPGVDYTITVYTWMSNYEDKSGSYYSPISINYRT | 2 |
| s31451 | TKLEVVAATPTSLLISWDFPAVLVTFYLITYGETGMNSPMQEFWVPGSM STATISGLKPGVDYTITVYAWYYERDEYQLIESSPISINYRT | 6 |
| s15487 | TKLEVVAATPTSLLISWDAPAVTVDFYLITYGEDGGNSPVQYFIVPGSVM TATISGLKPTVDYTITVYAYVVRSNFSFPSEISVYYSHGGG | 4 |
| s1137 | TKLEVVAATPTSLLISWDQPMWTVDFYVITYGEQGRNQPMQKFIVPGSV STATISGLKPWVDYTITVQAFHFRNSRSQPSPISMYGSGGGG | 6 |

| | | |
|---|---|---|
| s20460 | TKLEVVAATPTSLLISWDQPNVDVVFYLITYGETGGNKPVQEHWVPGEK STATISGLKPGVDYTITVYTYVRDGDLYADKPPISINYRTEG | 1 |
| s14335 | TKLEVVAATPTSLLISWDQSKVVVDSYVIAYGETHGNSPVQEFEVPGWT NTATISGLKPGVDYTITVYVFYVSAGDQMSGSPISINYRTSG | 5 |
| s22907 | TKLEVVAATPTSLLISWDAPSSNVDFYLITYGETHGNHPVQEHVVPGSM STATISGLKPGVDYTITVYTMYWGGSNGGYSYGHPISINYRT | 4 |
| s38496 | TKLEVVAATPTSLLISWDAPAVTVDYYVITYGERGGNSPVQEFIVPGSVR TATISGLKAGVDYTITVYAFVQHYDGHGSPYSSGHGSGGGS | 6 |
| s8891_s24482 | TKLEVVAATPTSLLISWDAPAGWVFFYRITYGETGGNSPVQESFVPGDM STATISGLKPGVDYTIYVYAQYDGSGEVGGFSFEPHSQYYRG | 3 |
| s20225 | TKLEVVAATPTSLLISWHAPNFYVDYYVITYGEHGYWPYYWQEDTVPGS MSTATISGLKPGVDYTITVYAGSNDDYNYYGSPISINYRTGG | 1 |
| s21007_s33549 | TKLEVVAATPTSLLISWDHPWVDQDLYFIVYGVTGGNSPFQEFTVPGSF STATISGLKPGVDYLITVYAQYYRDGWMYSHPISIFYRTGES | 3 |
| s5087 | TKLEVVAATPTSLLISWDMPALTVILYVITYGETGGNSPTQEFYVPGSKS TATISGLKPGVDYTITVYTVYEGDDETGYREDSPISINYRT | 1 |
| s7509 | TKLEVVAATPTSLLISWDMPAVSVDFYVITYGEVGGVHDSGTYEVLGWF STATISGLRGMVDYTITVYAHMVYDDNNSPHPISGFGSGSGK | 1 |
| s37107 | TKLEVVAATPTSLLISWMRPAVWVAFYVITYGEVGGNHPVQSFYVPGYV TTATISGLKPGVDYIITVYAWGTYSDEGFGSKSSGYGRAGGE | 6 |
| s22895 | TKLEVVAATPTSLLISWDFPMVTVDLYHIIYGEVFGTSPIQEFTVPGSKST ATISGLKPGVDYQIVVYAMVSYRDWYYMSPIYINYRTGLS | 4 |
| s6699_ds | TKLEVVAATPTSLLISWDMPAVFVFFYLICYGEIGGNSPMQCFRVPGSKS TATISGLKPGVDYVITVYTWYYSDSDGGATYSSPIIILYQT | 6 |
| s27652 | TKLEVVAATPTSLLISWWHPAVTVDFYVITYGETGGNSPTYRFIVPGSKN TATISGLSSDVDYTITVYAYMYGDGNKYPSPISINSGSGGG | 1 |
| s34966_s22269 | TKLEVVAATPTSLLIEWSTPMVTVDFYHITYGETGGNSPMQEFTVPGFV TRATISGLQPGVDYTITVYAYVSYIEYQYESPISINYRPGDG | 1 |
| s28784_s25968 | TKLEVVAATPTSLLISWDAPAVFVDFYLITYGETLGNSPVQEFFVPGSWS TATIEGLKPGVDYTITVYTWYYYDYHGLLYIYSPISINYRT | 1 |
| s1338 | TKLEVVAATPTSLLISWHFPMQTVDFYLIEYGETGGNSPVQVFTVPGST STATISGLKPGVDYTITVYAYYVEIVFYFPSPISINYRTGLK | 4 |
| s38112_ds | TKLEVVAATPTSLLISWDAPAVTVEFYLITYGEQGGNSPWQSFLVPGSV STATISGLKPGVDYTICVEALVARDGNHARWSGGCGSGGGGK | 1 |
| s36609_s25706 | TKLEVVAATPTSLLISWRHPAVTVQFYLITYGETGGNSPIQFHFVPGSKS TATISGLKPGVDYTITVYAYVYKYYPMFPSPISINYRTGPG | 1 |
| s37660 | TKLEVVAATPTSLLISWDHPWVTVVYYVITYGETGGNSPVQEFLVPGSV NTAEITGLKPGVDYTITVYATYYSSKKYEYVYSHPISINYRT | 1 |
| s31740_s38496 | TKLEVVAATPTSLLISWDKPAQTVDFYVITYGERGGNSPVQEFIVPGSVR TATISGLKAGVDYTITVYAFVQHYDGHGSPYSSGHGSGGGS | 1 |

| | | |
|---|---|---|
| s38285 | TKLEVVAATPTSLLISWDAPAVKVDYYVITYGEVGYWPYYWQEFTVPGS LSTAEISGLKPGVDYTITVYAGSYDQWYYWGKPISINYRTGG | 1 |
| s29734 | TKLEVVAATPTSLLISWDHPAVDVDLYHITYGETFGNSPMQSDTVPGSS RTATISGLKPGVDYTITVYAYYSSSDYYQPSPISINYRTGMK | 1 |
| s24419 | TKLEVVAATPTSLLISWDFPMVDVDFYRITYGETHENSPFQRDVVPGEK STATISGLKPGVDYTITVYAWYYGDSEKRIDTSHPISINYRT | 1 |
| s4171_s24471 | TKLEVVAATPTSLLISWDAPAVWVRFYEILYGETGGDSPYYWFTVPGSK STATISGLKPGVDYVITVYATYYASTRGWYRRVVPVVIDYRT | 1 |
| s36171_s28705 | TKLEVVAATPTSLLISWDAPALTVIFYVITYGETMGDSPLQQFVVPGSKS TATISGLKDGVDYLIIVYAYLHYSDDGYPPIISGGGGGGDG | 1 |
| s13507 | TKLEVVAATPTSLLISWDFPNDTYDFYFIQYGWGWFGNGPHQEFTVPG SMNTATISGLKPGVDYTITVYAYYGSGGFSPDPVQINYRTGGT | 1 |

**Table S2:** Cryo-EM data collection, refinement and validation statistics

| | #1 RBD-s19382 (EMDB EMD-72524) (PDB 9Y5Y) |
|---|---|
| **Data collection and processing** | |
| Magnification | 105,000 |
| Voltage (kV) | 300 |
| Electron exposure (e–/Å$^2$) | 42.5 |
| Defocus range (μm) | 0.8-2.8 |
| Pixel size (Å) | 0.84 |
| Symmetry imposed | C1 |
| Initial particle images (no.) | 5,071,075 |
| Final particle images (no.) | 230,942 |
| Map resolution (Å) | 3.16 |
| FSC threshold | 0.143 |
| Map resolution range (Å) | 2.0-41.0 |

| | |
|---|---|
| **Refinement** | |
| Initial model used (PDB code) | 7U0N |
| Model resolution (Å) | 3.31 |
| FSC threshold | 0.5 |
| Model resolution range (Å) | 2.0-41.0 |
| Map sharpening *B* factor (Å$^2$) | -80.6 |
| Model composition | |
| Non-hydrogen atoms | 11,837 |
| Protein residues | 1492 |
| Ligands | 0 |
| *B* factors (Å$^2$) | |
| Protein | 64.80 |
| Ligand | n/a |
| R.m.s. deviations | |
| Bond lengths (Å) | 0.004 |
| Bond angles (°) | 0.668 |
| Validation | |
| MolProbity score | 1.00 |
| Clashscore | 0.94 |
| Poor rotamers (%) | 0.00 |
| Ramachandran plot | |
| Favored (%) | 96.59 |
| Allowed (%) | 3.27 |
| Disallowed (%) | 0.14 |
